# A human embryonic stem cell model of Aβ-dependent chronic progressive neurodegeneration

**DOI:** 10.1101/419978

**Authors:** Teresa Ubina, Martha Magallanes, Saumya Srivastava, Charles Warden, Jiing-Kuan Yee, Paul M. Salvaterra

**Affiliations:** Department of Developmental and Stem Cell Biology, Beckman Research Institute of the City of Hope, 1500 E. Duarte Rd., Duarte, CA, 91010, USA; Integrated Genomics Core, Beckman Research Institute of the City of Hope, 1500 E. Duarte Rd., Duarte, CA, 91010, USA; Department of Diabetes, Beckman Research Institute of the City of Hope, 1500 E. Duarte Rd., Duarte, CA, 91010, USA; Biology Department, California State University San Bernardino, University Pkwy, San Bernardino, CA 92407, USA; The Irell and Manella Graduate School of Biological Sciences, Beckman Research Institute of the City of Hope, 1500 E. Duarte Rd., Duarte, CA 91010, USA

## Abstract

We describe construction and phenotypic analysis of a human embryonic stem cell model of progressive Aβ-dependent neurodegeneration (ND) with potential relevance to Alzheimer’s disease (AD). We modified one allele of the normal APP locus to directly express a secretory form of Aβ40 or Aβ42, eliminating the need for amyloidogenic APP proteolysis. Following neuronal differentiation edited cell lines specifically accumulate aggregated/oligomeric Aβ, exhibit a synaptic deficit and have an abnormal accumulation of endolysosomal vesicles. Edited cultures progress to a stage of overt ND. All phenotypes appear at earlier culture times for Aβ42 relative to Aβ40. Whole transcriptome RNA-Seq analysis identified 23 up and 70 down regulated genes (DEGs) with similar directional fold change but larger absolute values in the Aβ42 samples suggesting common underlying pathogenic mechanisms. Pathway/annotation analysis suggested that down regulation of extracellular matrix and cilia functions are significantly overrepresented. This cellular model could be useful for uncovering mechanisms directly linking Aβ to neuronal death and as a tool to screen for new therapeutic agents that slow or prevent human ND.

## Introduction

Alzheimer’s disease (AD) is a chronic progressive neurodegenerative disorder with a decade’s long preclinical phase. Clinical features include memory loss accompanied by progressive cognitive dysfunction, cortical atrophy and ultimately death. Neuropathology is well defined by a widespread accumulation of two prominent lesions in cortical brain regions: amyloid plaques and neurofibrillary tangles. Plaques are composed primarily of higher-ordered aggregates of small Aβ peptides derived from amyloidogenic proteolysis of a large transmembrane amyloid precursor protein (APP). Tangles are composed largely of hyperphosphorylated aggregates of a microtubule stabilizing protein tau, a product of the MAPT gene. All forms of AD exhibit accumulation of both Aβ plaques and neurofibrillary tangles and both are considered necessary for a definitive postmortem diagnosis [1]. Both plaque and tangle neuropathology correlate with decrements in cognitive function in AD, but our mechanistic understanding of how these lesions contribute to progressive neurodegeneration (ND) is still incomplete [2]. The reasons for this include the inherent complexity of the disease as well as inadequacies of current animal and cellular experimental models.

There are two general forms of AD: a rare autosomal dominant familial form (FAD) and the more prevalent sporadic form (SAD). Both FAD and SAD have a complex genetic component (i.e. >20 identified risk alleles with APOE4 being the most prominent), significant life-style associations (obesity, sleep, exercise, etc.), several associated co-morbidities (i.e. diabetes, head trauma), and of course the most important correlate of all, old age. This complexity coupled with the lengthy time course of the disease process and the relative inaccessibility of patient samples makes it challenging to develop effective therapies. Most clinical trials of AD therapeutics have failed at an unprecedented rate [3]. Part of this dismal status quo is likely due to our poor understanding of basic pathobiology of AD. Preclinical testing of AD drugs is typically done using transgenic mice, but these AD models are deficient in at least two important ways. They fail to exhibit progressive neurodegeneration (ND) and do not generally display neurofibrillary tangle pathology [4].

Transgenic rodent AD models (see for example: https://www.alzforum.org/research-models/alzheimers-disease) usually rely on over-expression of one or more FAD mutant genes. They generally exhibit cognitive phenotypes, synaptic deficits, and accumulation of aggregated Aβ through amyloidogenic proteolysis but not progressive ND or tau related pathology suggesting their use as preclinical models is not practical [5]. Progressive ND and tau related pathology are also not generally observed in aged non-human primate models [6]. Considerable effort has gone into developing mouse models with tau AD-like tau pathology driven in large part by notable differences in human and mouse tau isoforms differences in tau [7] or incorporating a mutant tau MAPT gene [8]. MAPT mutations, however, not associated with AD can cause other types of non-AD ND disease [9]. Generation of “second generation” knock-out/knock-in rodent models, designed in part to eliminate over-expression artifacts common with high level transgene expression and humanizing the APP sequence also lack progressive ND or tangle pathology but have been useful in highlighting interpretive phenotypic complexity of “first generation” models [4].

A few distinctive AD models have been developed which exhibit progressive Aβ-dependent ND using direct over-expression of secretory Aβ coding transgenes. This approach eliminates the need for amyloidogenic proteolysis of APP to generate Aβ. These types of models have been well characterized in invertebrate organisms such as *Drosophila* [10] and less extensively in mice [11–14]. Direct Aβ over-expression in rodent or *Drosophila* neurons does not significantly affect normal brain development but results in an impressive range of putative AD-like phenotypes including accumulation of aggregated/oligomeric Aβ42, neurological and memory deficits, early and massive autophagy/endosomal/lysosomal abnormalities, mitochondria dysfunction and plaque like accumulations of Aβ [11,13–17]. None of these models exhibits tau pathology and they all suggest that phenotypes are exclusive to Aβ42 overexpression since an equivalent amount of Aβ40 does not result in similar changes. Direct expression of Aβ may thus be an effective way to study the now uncharacterized cascade of Aβ–dependent mechanistic changes which are thought to precede ND as postulated in the dominant amyloid cascade hypothesis [18]. While this hypothesis is strongly supported by a wealth of supporting evidence [19], it remains “controversial” because of seemingly discordant observations made in phenotypically deficient animal models or more importantly correlative clinical discrepancies in AD patients [20]. The most serious discrepancy often cited is the relative timing of amyloid and tangle pathology with respect to cognitive status of patients, however, recent longitudinal imaging studies have generated a better time line for pathogenic progression in patients and convincingly place Aβ at the beginning, at least in FAD patients [21].

Explanations for phenotypic deficiencies in current AD models are often attributed to the relatively short lifespan of rodents or human species-specific factors. These human differences are either known (i.e. isoform differences in tau or Aβ sequence, etc.) or unknown, but possibly related to the complex genetic context of both FAD and SAD and thus not possible to adequately model in non-human animals. With the advent of reprograming technology, AD patient derived iPS culture models have been established which now allow phenotypic characterization in a human genetic context [22]. Pioneering studies document a number of promising AD-relevant phenotypes using cells derived from with FAD patients including increased amyloidogenic Aβ production/aggregation, increased ratios of Aβ42/Aβ40, or lysosomal/endosomal dysfunction [23,24]. The AD-relevant phenotypic repertoire has even been extended to include tau related pathology when FAD genes are overexpressed in human neural precursor cells or FAD iPS cells are differentiated in a 3-D culture format [25,26]. The authors suggest that the AD relevant tau phenotypic extension could be due to the more complex “brain-like” cellular organization of 3-D cultures and/or decreased removal of extracellular Aβ during normal culture media replacement [27]. Progressive ND, however, was not among the phenotypes of these 3-D cultures. Current human iPS models while encouraging are still not suitable to investigate mechanisms linking Aβ to progressive ND.

Here, we describe the construction and initial AD-like phenotypic characterization of a new human embryonic stem cell model Aβ-dependent ND. We used genomic editing to modify parental WiCell WA09 cells (H9) to directly express a secretory form of either Aβ40 or Aβ42 from one allele of the normal APP gene locus. Expression is thus under control of the normal APP promoter, but amyloidogenic processing is not necessary for Aβ production. Following neuronal differentiation, edited neurons accumulate intracellular aggregated/oligomeric Aβ but the rate is faster for Aβ42 edited lines. Aggregated/oligomeric Aβ preferentially localizes near fragmented/pyknotic nuclei, even in unedited cells which presumably produce a small amount through amyloidogenic processing. Aβ42 edited cells elaborate several other AD-relevant phenotypes at a faster rate than Aβ40 lines including synaptic deficits and a greater accumulation of endolysosomal vesicles. Importantly, both edited genotypes exhibit progressive ND relative to unedited control cells and the rate of progression is faster for Aβ42 cultures. Whole transcriptome RNA-Seq analysis identified a small set of differentially expressed genes (DEGs) in the Aβ42 samples compared to unedited samples which had a similar directional fold-change in Aβ40 samples but a smaller magnitude. This suggests that common genetic pathways may be affected since mRNA was isolated at a time when phenotypic changes were more extensive or exclusive to Aβ42 samples. Functional annotation and pathway analysis of DEGs identified “increased neuronal cell death” and “decreased memory” as the highest and lowest scoring functions perturbed in Aβ42 edited cells and suggested that disruption of extracellular matrix and cilia play a prominent role.

## Methods

### Genomic Editing

TALEN (Transcription activator-like effector nuclease) pairs were designed to target DNA upstream of the normal APP translation start site using published criteria, their cutting efficiency established in HEK293T cells and used to generate a double strand break in the APP target [28]. Donor templates for homology repair contained homology arms flanking the targeted site along with a secretory signal derived from the rat proenkephalin (PENK) gene, a human Aβ40 or Aβ42 coding sequence, and a polyA tail. Donor templates also contained a puromycin selection gene under control of the human phosphoglycerate kinase.

H9 (WiCell WA09) human embryonic stem cells were obtained from the WiCell Foundation and cultured on a feeder free system (Matrigel). Cells were harvested at appropriate confluency and nucleofected with TALEN pairs and donor template using an Amaxa Nucelofector. Nucleofected cells were grown for 48 hours, harvested and plated on puromycin resistant feeder cells at a dilution of 1/30 for 48 hours and then transferred to puromycin drug selection media for two weeks. Approximately 1/2 of appropriate size colonies were collected for PCR analysis using primer pairs that spanned the flanking DNA and the donor plasmid sequences to confirm insertion of the expression cassette. The stem cell colonies positive for correct size PCR fragments at both the 3’ and 5’ sites were expanded and analyzed for expression of edit specific Aβ40 or Aβ42 expression using qRT-PCR analysis. The forward primer was specific to the rat secretory signal sequence (not present in the human genome) and the reverse primer targets the end of the Aβ40 sequence. The specific sequences and editing and verification details are included in the **Supplemental Methods and Data**.

### Cell culture

ESC culture, embryoid body generation and neuronal differentiation were adapted from a well-established protocol [29]. Briefly, stem cells were grown in gelatin coated six-well plates on an irradiated mouse embryonic fibroblasts feeder layer. Stem cells were maintained in HuES medium which was replaced daily and differentiating colonies were manually removed to maintain pluripotency. Stem cells were passaged weekly and differentiation was initiated ~1 week after passage using dissociated cells transferred to a 10 cm culture plate for embryoid body (EB) generation. On day 3 cells were grown in Neural Induction Media (NIM) with N2 supplement and 2 µg/ml heparin. On day 5 media was supplemented with ascorbic acid, trans-retinoic acid, Y-27632 ROCK inhibitor, and BDNF. On day 7 smoothened agonist 1.3 was added. Media was replaced every 3^rd^ day and after ~28-31 days EBs were collected, rinsed with Ca^2+^/Mg^2+^free PBS, dissociated into individual cells and plated in either 6 or 24 well culture plates precoated with poly-L-ornithine and laminin (1.7×10^6^ or 0.34×10^6^ cells per well) in neural differentiation medium supplemented with 25 μM β-mercaptoethanol and 25 μM glutamate. Cultures were initially treated with 0.5 µM ethynyl deoxyuridine (EdU) for 24 hrs and weekly thereafter up to ~50 days to maintain only post mitotic cells. Complete media recipes, suppliers and protocol details are included in the **Supplemental Methods and Data**.

### qRT-PCR

Total RNA was extracted using the RNeasy Micro Kit from (Qiagen) following the manufacturer’s protocol. RNA concentration and purity was determined spectrophotometrically and cDNA prepared using qScript cDNA SuperMix (Quanta) following the manufacturers protocol. All reactions were carried out in a 20 µl reaction mixture containing 12.5 µl iQTM SYBR® Green Supermix (Bio-Rad), 2 µM of each forward and reverse primer, 0.25 µg cDNA, and DEPC-Treated Water (Ambion) to adjust the final volume to 20µL. Amplification was carried out using a BioRad CFX96 Touch^™^ Real-Time PCR machine in clear 96 well sealed plates and data was collected and analyzed using BioRad CFX Manager (v3.1). Additional details and primer sequences are included in the **Supplemental Methods and Data**.

### Microscopy, Immunocytochemistry, Live-Dead Analysis and Image Analysis

Fluorescence samples were observed with a Zeiss Axio Observer microscope (Xenon illumination) using either a 20X NA=0.80 plan-apochromat objective or a 40x or 63x plan-apochromat objective (NA=1.4, Oil). Optical Z sections were acquired with a Zeiss Axiocam506 camera using Zeiss Zen Blue microscope control software (SP2). Unstained cultures were observed using a Nikon Diaphot inverted microscope equipped with Hoffman modulation contrast objectives (HMC EF 10X NA=0.25 or HMC 20X LWD NA=0.4) and images were obtained with a SPOT RT230 cooled CCD camera operated by SPOT Advanced Imaging Software. Image analysis used semi or fully automated macros implemented in the FIJI version of NIH ImageJ (v1.46 or 2) [30]. For visual clarity some images are adjusted for brightness and contrast using Adobe Photoshop (CS4 or CS5). Due to variability in the number of cells in neuronal clusters both among genotypes differentiated in parallel, as well as across independent differentiations, quantitative data were usually normalized to the number or area of DAPI staining.

### Antibody staining

Cells were grown on polyornithine/laminin coated 15mm No.1 glass coverslips (Fisher Scientific) placed in 6 well plates. Cells were fixed with 4% paraformaldehyde for 20 minutes followed by washing in PBS (3x, 5 min.) and coverslips were stored in 0.03% NaN_3_ in PBS at 4°C until observation. Coverslips were incubated with blocking buffer (0.3% Triton X-100 and 5% Bovine Serum Albumin in PBS) for ~2 hr. at room temperature, washed briefly with PBS and incubated overnight at 4°C with primary antibody diluted in 0.3% Triton X-100, 1% bovine serum albumin in PBS (antibody dilution buffer). Coverslips were washed with PBS (3×5 min.) with antibody dilution buffer and incubated with fluorescent labeled secondary antibodies for two hours at room temperature, washed with PBS, incubated with DAPI (1µg/µl) for 5 minutes at room temperature, washed with PBS (2x, 5 min.) and mounted onto glass slides using DAKO Fluorescent Mounting Medium. Additional coverslips were stained after eliminating either the primary or secondary antibody to serve as negative staining controls. Specific antibody staining details and image analysis parameters are included in the **Supplemental Methods and Data**.

### Live-Dead Analysis

Neuronal viability was estimated by measuring the relative proportion of live/dead cells in neuronal clusters grown on coverslips or directly in culture wells using a commercial fluorescence assay (ThermoFisher LIVE/DEAD^™^ Viability/Cytotoxicity Kit, for mammalian cells, #L322) according to the manufacturer’s directions. Additional details and image analysis parameters are included in the **Supplemental Methods and Data**.

### Statistical Analysis

We used Prism (v7, Graph Pad) for statistical analyses (descriptive statistics, ANOVA, variance estimates and correlation) and graphic preparation.

### RNA-Seq

Stem cells were differentiated for 36 or 38-days and total RNA was extracted using the RNeasy Micro Kit (Qiagen) following the manufacturer’s protocol. RNA concentration and purity were determined using a NanoDrop ND-1000 spectrophotometer and processed for RNA-Seq analysis by the City of Hope Genomic Core Facility. Detailed processing and analysis protocols are included in the **Supplemental Methods and Data**. The sequencing data files have been deposited in the NIH GEO database (GSE119527).

## Results

### Model construction

We used TALEN genomic editing to modify the normal wild-type APP gene in WiCell WA09 (H9) human embryonic stem cells (hES). This cell line was chosen because of its widespread use in stem cell studies, the availability of many well characterized neuronal differentiation protocols and because the APOE genotype contains one copy of an ε4 allele which is the major genetic risk factor for SAD [31]. The APOE genotype (ε4/ε3) was confirmed using allele specific PCR analysis (not shown). The editing strategy is shown schematically in Fig. 1. TALEN pairs were designed to induce a double strand break (DSB) within the first exon of the *App* locus upstream of the normal *App* transcriptional start site. The DSB was repaired by homologous recombination in the presence of donor plasmids that contained a secretory signal sequence derived from the rat preproenkephalin gene (PENK, *Rattus norvegicus*) fused in frame to either a human Aβ40 or Aβ42 coding sequence and followed by a polyA tail just upstream of a puromycin drug selection gene. This insertion cassette was flanked by left and right homology arms to direct insertion into the normal *App* locus.

**Fig. 1.**
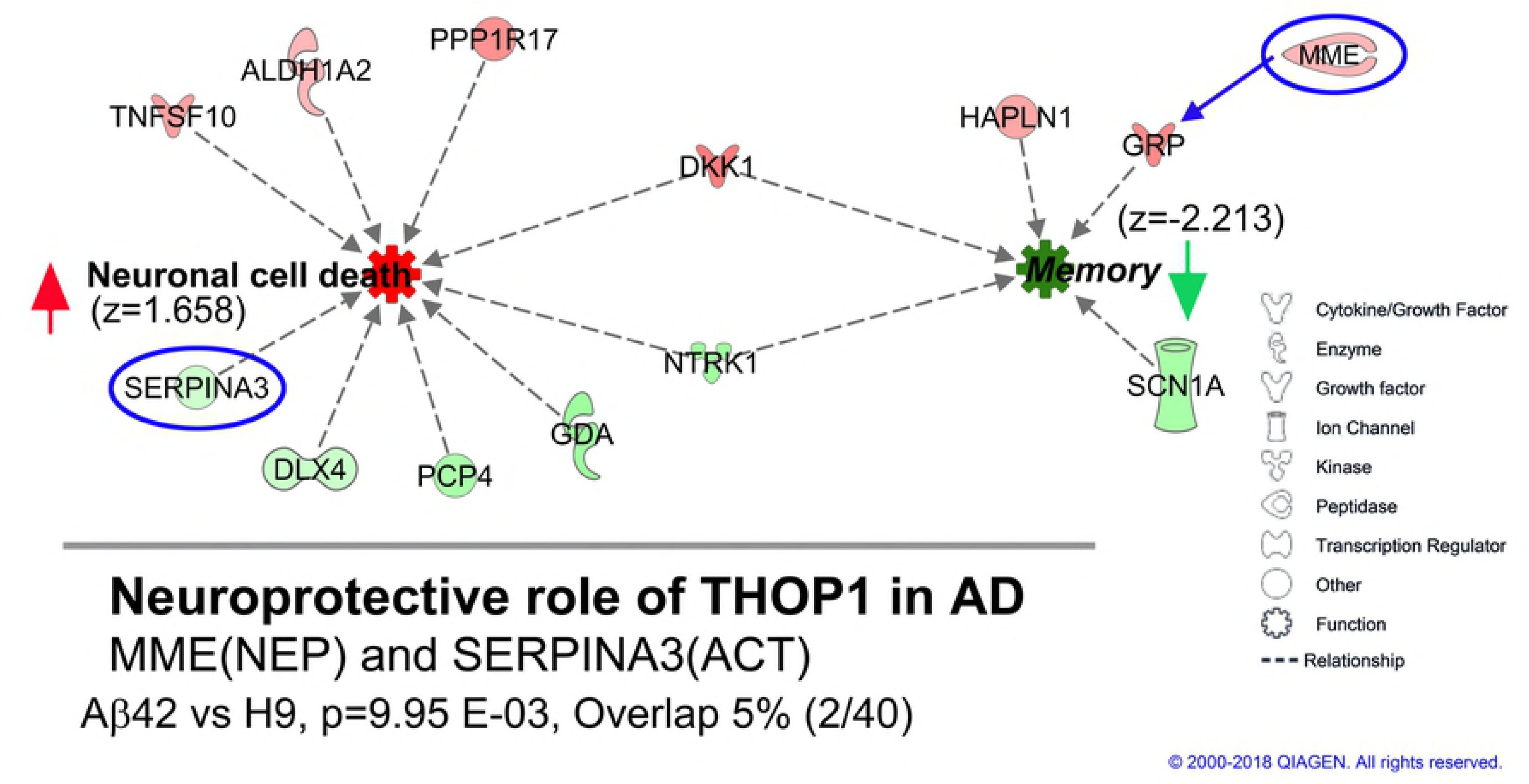
Genomic editing of APP gene locus. TALEN pairs were designed to target and induce a double strand break (DSB) in the first exon upstream of the normal APP translation initiation codon (APP ATG). The DSB was repaired by homologous recombination in the presence of plasmids containing the coding sequence for either Aβ40 or Aβ42 fused in frame with a rat preproenkephalin secretory signal sequence (SS) and followed by a polyA tail (not shown). Repair plasmids additionally included a PGK puromycin drug selection gene (Puro) and were flanked by left and right homology arms homologous to APP flanking sequences (HAL, HAR). Cassette insertions were confirmed by genomic PCR using specific primers in either the HAL (5’) or the HAR (3’) and a site in the insertion cassette. This editing strategy simultaneously inactivates one APP allele and replaces it with a cassette that directly expresses a secretory form of either Aβ40 or Aβ42 under normal APP regulatory control. The specific sequences and other details are included in the **Supplemental Methods and Data.**

Successful editing resulted in inactivation of the modified *App* allele and its replacement with direct expression of either secretory Aβ40 or Aβ42 Importantly, the parental and edited cell lines are essentially isogenic ensuring that phenotypic differences are directly attributable to the specific edits. The rat PENK secretory signal sequence is not present in the human genome allowing PCR analysis to specifically detect edited Aβ transcripts. Following translation, the signal peptide is completely removed by normal secretory pathway processing resulting in direct production of either an Aβ or Aβ42 peptide [11,15] eliminating any requirement for amyloidogenic APP processing by β and γ secretase. Since the edits are introduced directly into the normal APP locus, expression will be under control of the normal APP regulatory DNA. This distinguishes our model from others that generally used exogenous promoters to drive overexpression. We hypothesized that this model could potentially speed up proteotoxic Aβ accumulation on a time scale suitable for working with cultured human neurons while potentially minimizing overexpression artifacts.

Proper editing was initially identified by PCR screening of multiple subclones using 3’ and 5’ specific primers and confirmed by genomic sequencing. Since subcloning as well as TALEN editing has the potential to generate off-target effects (primarily indels) or other mutations, although at extremely low levels [32], we phenotypically characterized two independently isolated subclones for each edited genotype in parallel. We noted no consistent phenotypic differences between subclones suggesting that the differences we describe are genotype specific (i.e. due to direct expression of either Aβ40 or Aβ42). All edited cell lines used in this study were heterozygous for the edit ensuring that normal APP will still be expressed from the unedited allele.

### Aβ and APP expression

We used qRT-PCR to measure edit specific expression of secretory Aβ using a forward primer specific to the rat PENK secretory signal peptide which is absent from the human genome and a reverse primer to the end of the Aβ40 sequence which is present in both edits. As expected, no edit specific transcripts were detected in unedited H9 cells (Fig. 2A). Significant levels were found in undifferentiated stem cells, EB stage cells or differentiated neurons. The relative expression levels were similar for both edited genotypes at these three developmental stages indicating that they are under the same regulatory control. We additionally confirmed that only secretory Aβ42 expression could be detected in Aβ42 edited lines using a reverse primer specific to the unique 5’ nucleotides in Aβ42 (not shown). Undifferentiated stem cells show an intermediate expression level, consistent with normal APP expression previously reported at this stage [33]. Transcript abundance decreased significantly during EB formation and increased to the highest levels in 10-day old neuronally differentiated cultures. The relative ratio of edit specific Aβ mRNA for stem cells, embryoid bodies and differentiated neurons was ~20:1:100. We expect that Aβ protein levels would likely be highest in differentiated neurons (i.e. ~ 5-fold greater than in stem cells).

**Fig. 2.**
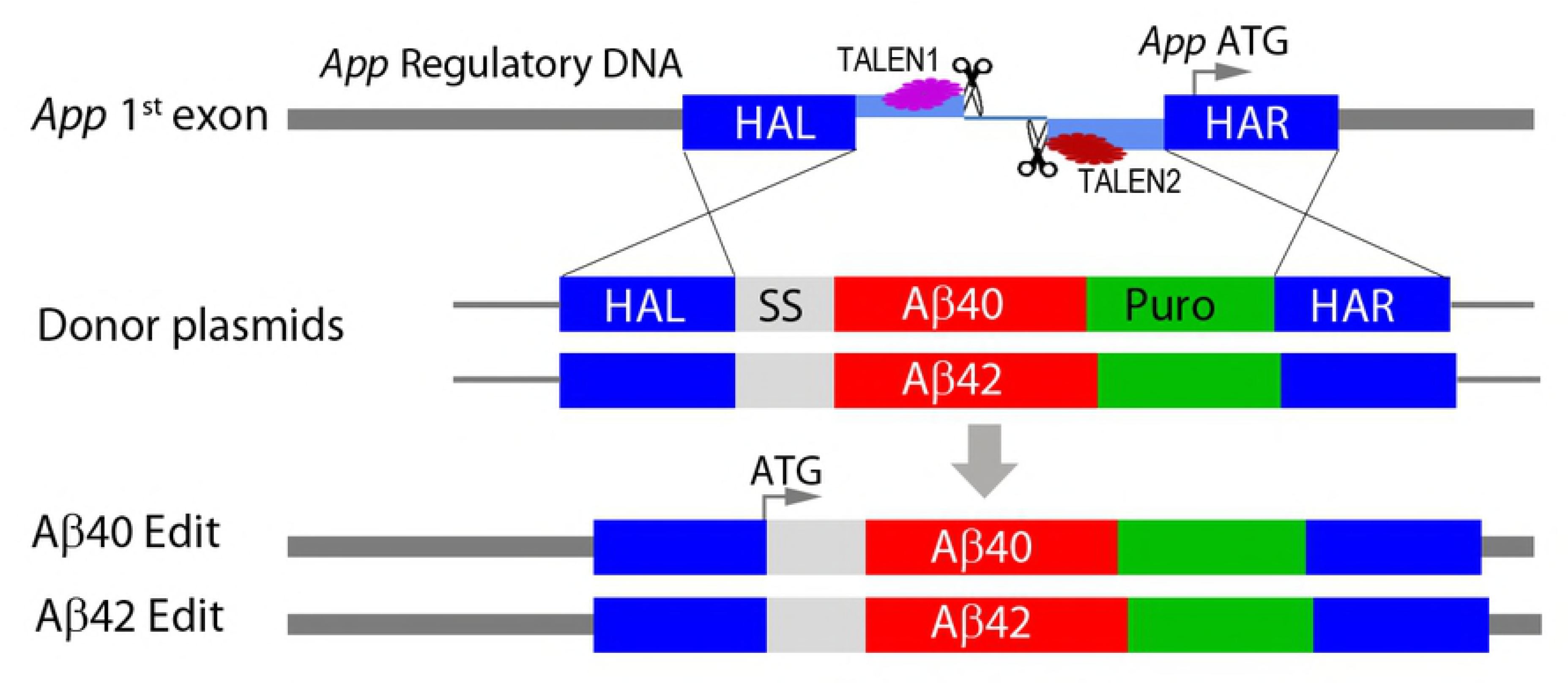
(A) Direct expression levels of edit specific Aβ are similar for both edited genotypes and dynamic during early stages of differentiation. Stem cell cultures have intermediate expression, embryoid bodies have significantly less expression and differentiated neurons have the highest, reaching maximal expression by ~10-20 days after EB dissociation and plating in neural differentiation medium. The relative ratios of Aβ expression were ~1: 0.05: 5 for the 3 developmental stages. There were no significant differences in expression level comparing edit specific Aβ40 and Aβ42 at any stage (ANOVA, Dunnett’s correction). No significant secretory Aβ expression was detected in unedited H9 samples. Data were from 6 independent stem cell cultures, 4 EB stage cultures and 22 individual 10-20 day old differentiated neuronal cultures. **(B) Editing does not affect APP expression from unedited alleles**. We used primer pairs spanning 3 different APP exons. The pattern of expression was similar for all 3 genotypes and average relative expression for the primer pairs was 1: 0.71: 0.5 for H9: Aβ40: Aβ42 and is consistent with expected inactivation of one APP due to editing. Expression of edit specific Aβ was ~30-fold less than APP expression and is replotted from (**A**) for comparison. Data were from 4 independent differentiations of H9 cells and 8-20 differentiations for edited genotypes taken from 10-34-day old cultures. In (**A**) expression was measured by qRT-PCR using a forward primer specific to the secretory signal sequence (not present in the human genome) and reverse primer to a sequence common to Aβ40 and Aβ42. In (**B**) the forward and reverse primers spanned indicated exons in the APP sequence. Bars are mean normalized expression (MNE) relative to GAPDH (±STD).

We additionally measured APP expression in 10-day old differentiated neurons using forward and reverse primers that span different adjacent exons along the length of the normal neuronal APP transcript (Fig. 2B). Different exon spanning primer pairs detected APP transcripts over an approximately ~8-fold range, but the pattern was similar for all 3 genotypes. The average relative APP expression for all 3 primer pairs compared to H9 was 0.71 for Aβ40 and 0.5 for Aβ42 a result is consistent with expected inactivation of only the edited APP allele. This confirms that editing does drastically affect APP expression from the unedited allele.Unexpectedly, however, direct Aβ expression was ~30-fold lower relative to APP expression (the Aβ data is replotted from Fig. 2A). This could be due to weakening of a regulatory element in the first intron of APP [34] or alternatively to negative interference of the drug selection gene present in the insertion cassette [35]. Whatever the reason, direct expression levels for edit specific Aβ are significantly lower than APP.

Unfortunately, we were unable to reliably measure Aβ protein levels in either immunoprecipitated culture supernatants (10 ml of immunoprecipitated sample pooled from 5 samples every 2 days from a single well of a 12 well culture plate), or in guanidine hydrochloride or formic acid cell extracts (prepared from 2 individual 12 well cultures) using commercial ELISA assay kits (Invitrogen, Aβ40 #KHB3481, sensitivity 6 pg/ml; Aβ42 #KHB3441, sensitivity = 10 pg/ml). These negative results are consistent with our qRT-PCR analysis and suggest that Aβ peptide levels in our cultures are significantly lower than those generated by amyloidogenic APP processing in differentiated neuronal culture models derived from human FAD iPS cells or cells transduced with FAD genes [25,36,37].

### Early development and culture differentiation

AD is a chronic and progressive neurodegenerative disease that only appears later in life. We observed no consistent genotype specific differences in morphology of ES stage culture, embryoid body (EB) formation or the earliest stages of culture in neuronal differentiation medium (see **Supplemental Methods and Data, Fig. S1**). Additionally, earlier stage embryoid bodies (7 day old) lose their initial positive staining for OCT4 (stem cell marker) and acquire Nestin staining (early neural differentiation marker) at a similar time independent of editing (see Supplemental Fig. S1). The appearance of differentiation markers in 10-day old cultures is shown in Fig. 3. The total cell number (DAPI), DCX positive cells (doublecortin, early stage neuronal differentiation) and NeuN positive cells were not significantly different among the tree genotypes (ANOVA, Dunnett correction). We conclude that genomic editing and APP heterozygosity do not appear to affect neurogenesis or early neural development in our cultures and that the majority of cells (60-70%) can be classified as neurons after 10 days. Hereafter all culture ages for differentiated cells are specified relative to EB dissociation and plating taken as day 0.

**Fig. 3.**
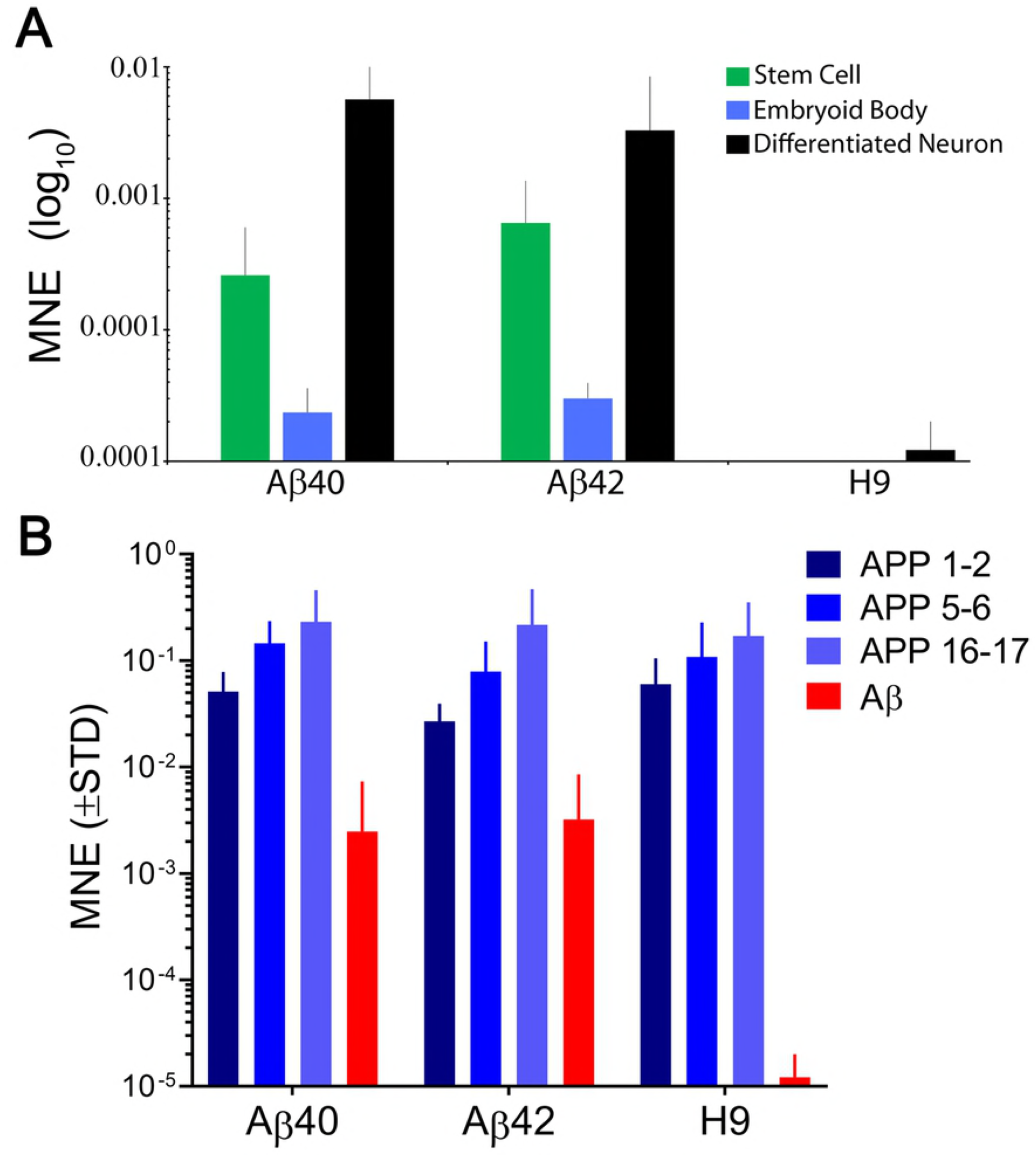
Editing does not significantly affect early stage neuronal differentiation. **(Left)** Representative images of 10-day old cultures stained with antibodies to DCX (doublecortin, green) to visualize early stage neuronal differentiation, NeuN (red) to visualize more mature neurons and DAPI (blue) to assess total cell number. **(Right)** Quantification of positively stained cells for each marker indicate that there were no genotype specific differences (ANOVA, Dunnett’s correction). Bars are the mean (SEM) of 3 biological replicates. Scale bar = 30 µm.

Consistent with the neuronal maker data the morphological appearance of all three genotypes, as well as the independent edited clones, remains quite similar up to about 30-days of culture (Fig. 4). One day old cultures have only isolated cells, a few of which appear to exhibit short processes. By ~15 days, cells appear to self-organize into loosely defined neural clusters (NC) and elaborate neural processes, some connecting to adjacent clusters. The size of the NCs increases slightly between 20 and 30 days and begins to appear more 3-D dimensional. Many NCs are connected to each other by neural processes at this stage. The size of NCs in both edited genotypes often appeared slightly larger compared to H9 cultures, but this was not statistically significant (ANOVA, Dunnett corrected) and absent by 40 days.

**Fig. 4.**
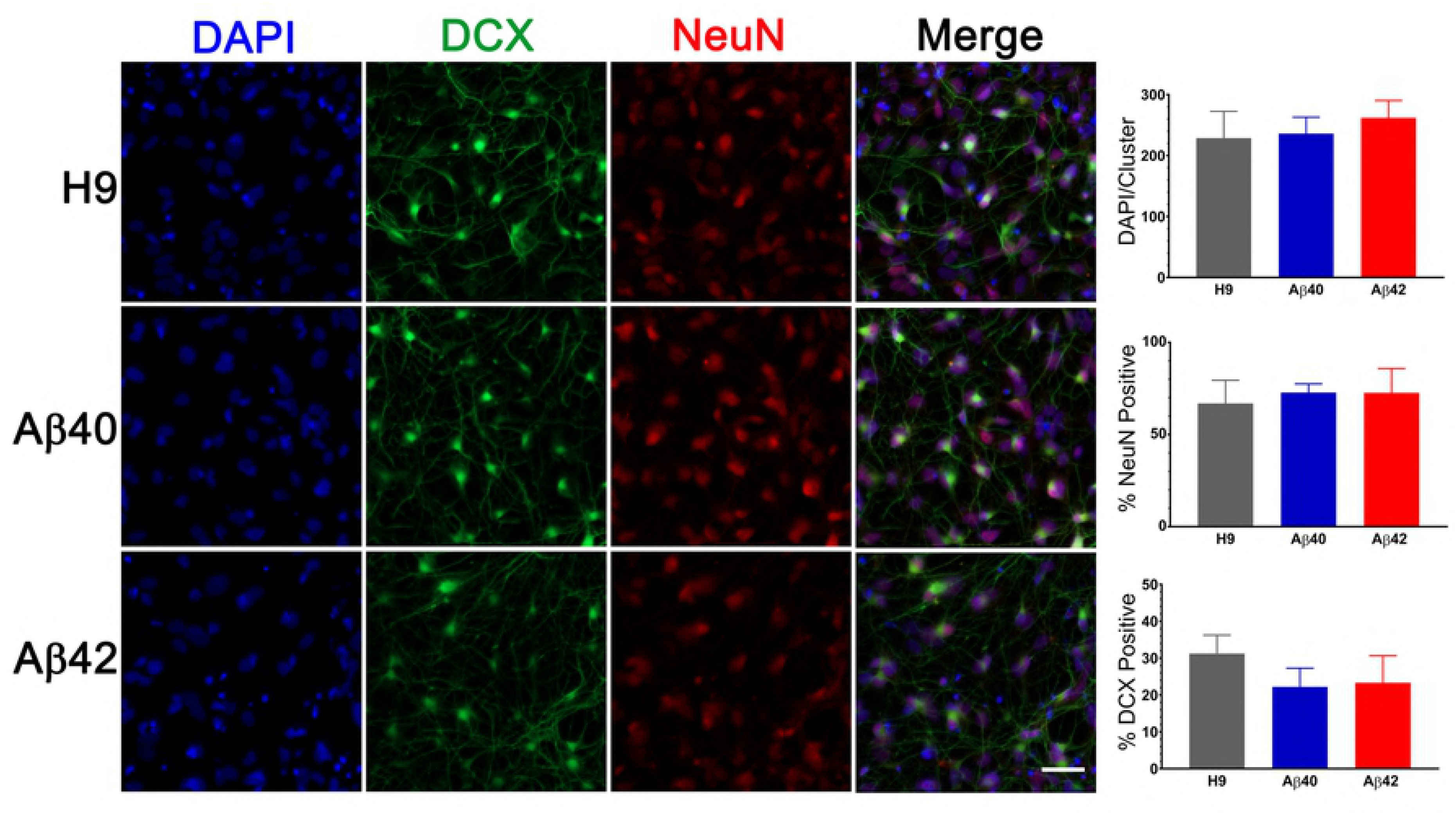
Representative Hoffman interference contrast images of unedited H9 parental cells and two independently isolated clones for each edited genotype (Aβ40:#31, #41 and Aβ42:#14, #26) at different culture ages. Isolated cells in 1-day cultures begin to cluster together a few days after plating. By ~10-15 days of differentiation all 3 genotypes form more recognizable neuronal clusters (NC) which are attached to the culture surface and elaborate neural processes which connect with adjacent NCs. Morphologic appearance of all 3 genotypes was generally similar up to ~30-40 days of culture. The absolute size of NCs varied across independent differentiations, however, there were no significant differences among the 3 genotypes up to ~30 days of age (ANOVA, Dunnett correction). After ~20-30 days, Aβ42 genotypes begin to exhibit a granular and darker appearance (especially evident in the Aβ42 clone #26 30 day image) and the somal regions are no longer firmly attached to the culture surface but tethered by their neuronal processes. After 50-60 days, essentially all Aβ42 genotypes exhibit this type of morphology as do many of the Aβ40 cultures at culture times greater than ~70-90 days. We did not observe any consistent clone specific differences for edited genotypes. Scale bars = 10 μm for 1-day culture and 100 μm for other ages.

At culture times of 40-50 days, Aβ42 NCs usually had a more granular appearance and were darker than the other genotypes. In one case we also observed this morphologic change as early as 30 day (see Fig. 4, Aβ42 clone #26). This morphologic appearance was more prominent in Aβ42 NCs older than 60 days and thus appears to specific to the Aβ42 edited cells. The neuronal soma for both edited genotypes lost firm attachment after ~60-70 days but still remained loosely tethered to the culture dish through their neural processes. This could be easily observed when gently moving the culture dish and was never seen in the unedited H9 cultures. Notably, we were not able to culture viable cells for either edited genotype for any time longer than 120 days. In contrast, unedited H9 cultures could be maintained for >266 days. Editing thus decreases the survival time of neurons and results in specific morphologic changes, especially apparent for Aβ42 edits. The absolute size of NCs had considerable variation in independent differentiations but this was a property of all 3 genotypes. These morphologic descriptions were generalized from observations made by 3 different investigators on 15 independent differentiations over a period of >2 years using several different lots of media and supplements.

### Alzheimer’s related phenotypes

#### Accumulation of aggregated/oligomeric Aβ and pyknotic nuclei

The main objectives of this study were to document putative AD-related phenotypes resulting from direct Aβ expression in human neurons and to compare the extent of phenotypic differences between Aβ40 and Aβ42. The most commonly observed AD-related phenotypes present in most animal models as well as several iPS culture models is the accumulation of aggregated Aβ produced by amyloidogenic APP proteolysis (see [4,22] for reviews).

We double stained cultures with an anti-Aβ antibody (7A1a) which specifically recognizes low and high molecular weight aggregates/oligomers of Aβ40 or Aβ42 [17,38] and anti-Tuj1 (TUBB3 gene product) to confirm neuronal cellular identity. In 32 day old cultures the level of 7A1a positive staining is genotype specific (Fig. 5A). The relative area of 7A1a staining (normalized to Tuj1) was minimal in H9, intermediate in Aβ40 and significantly higher in Aβ42 cultures. Compared to unedited H9 cultures, the area of 7A1a staining was ~2 fold higher in Aβ40 cultures (but not statistically different from H9) and ~3 fold higher in Aβ42 cultures (p<0.0016) at 32 days (Fig. 5B). At a later culture age (63 d) the average accumulation of 7A1a positive staining relative to H9 increased to ~3 fold in Aβ40 and ~4.5 fold in Aβ42 cultures. Accumulation of aggregated/oligomeric Aβ is thus progressive and faster for Aβ42 relative to Aβ40 cultures. This result is consistent with the biophysical aggregation properties of these 2 peptides *in vitro* [39] and since both edited genes are expressed at comparable levels suggests that Aβ42 may be removed at a slower rate. Both edited genotypes have less Tuj1 positive staining which was especially evident in older cultures but not in older H9 cultures (5A).

**Fig. 5.**
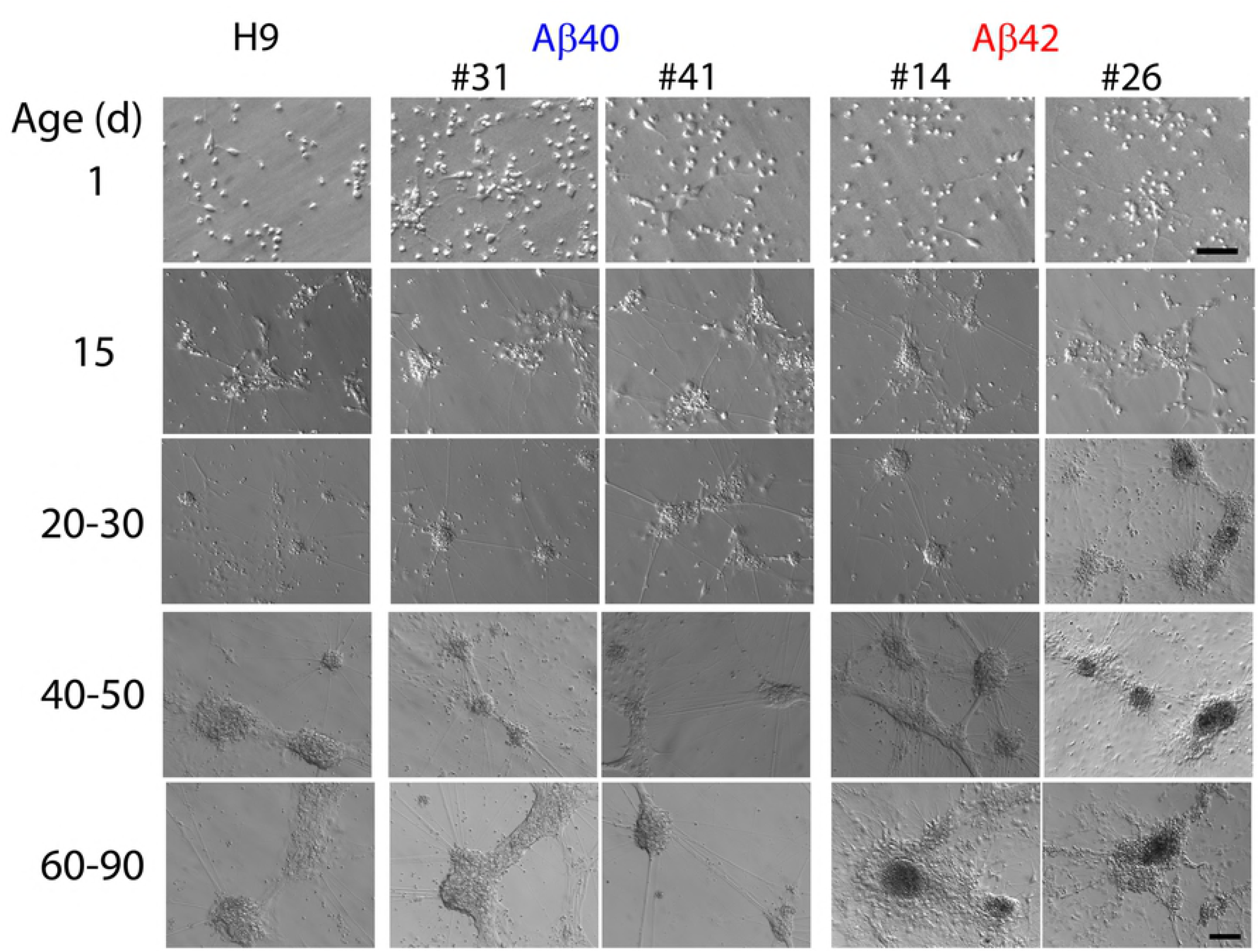
Accumulation of aggregated/oligomeric Aβ is time dependent and more prominent in Aβ42 relative to Aβ40 edited cultures and is associated with pyknotic nuclei, even in unedited H9 samples. (**A)** Maximum intensity Z-projections of NCs fluorescently stained with anti-Tuj1 (neuronal, green) and anti-Aβ 7A1a (aggregated/oligomeric Aβ, red) antibodies in 32 or 63 day old cultures. Consistently, the area of 7A1a positive staining is greater in Aβ42 NCs, intermediate in Aβ40 NCs and much lower in unedited H9 cultures. Staining is primarily intracellular and initially appears as small puncta which are more obvious in areas of lower staining intensity. (**B)** Box and whisker plot of relative 7A1a staining in individual NCs (normalized to Tuj1 staining). The line in the box is the median value, whiskers are the range. Data is from 4 independent differentiations. NCs from Aβ42 cultures have significantly greater accumulation of aggregated/oligomeric Aβ at 32-days (ANOVA, Dunnett correction) relative to H9. Accumulation in Aβ40 cultures appears higher than H9 but are not significant at this age. Mean relative accumulation of 7A1a staining ±SEM were: H9 = 1±0.235, Aβ40 = 3.77±0.704, Aβ42 = 6.93±1.63. In 63-day old cultures, both Aβ40 and Aβ42 are significantly different relative to H9. The mean relative areas are: H9 = 1±0.157, Aβ40 = 2.34±0.287, Aβ42 = 3.959±0.337). (**C)** 7A1a staining is present primarily in areas near pyknotic/fragmented DAPI stained nuclei (i.e. small intensely fluorescent structures, arrowheads) and absent from cells with normal nuclei (i.e. large, weak DAPI fluorescence, arrows). Images are from a single optical section of a 32-day old Aβ42 sample (top row) with a magnified view (bottom row) of the indicated rectangular area. (**D)** Association of 7A1a and pyknotic nuclei is not dependent on editing. (**Left**), images of fragmented or intact nuclei from Aβ42 or H9 cultures. (**Right**), spatial distribution of 7A1a fluorescence relative to the center of mass for DAPI staining. Bars are the mean (SEM) area of 7A1a staining in individual concentric circles centered on the DAPI staining. Data is from at least 60 nuclei or pyknotic nuclei from 3 independent differentiations of 32/34 day old cultures. Scale bar in **A** = 10 μm, **C**= 20 μm, **D** = 4 μm.

7A1a staining was primarily intracellular and appeared to be in close proximity to pyknotic nuclei characteristic of dead or dying cells (i.e. nuclear condensation and fragmentation). Normal neuronal nuclei are large and only weakly stained with DAPI while pyknotic bodies are smaller and have intense DAPI fluorescence. Fig. 5C shows this spatial relationship in a 32-day old Aβ42 culture. Larger areas of 7A1a staining were generally absent in areas near normal nuclei but common near pyknotic nuclei. Whenever 7A1a staining was occasionally present close to normal nuclei the staining area was small and punctate (possibly vesicular).

We also noticed that the few cells in unedited H9 cultures with 7A1a positive staining also seemed to be near pyknotic nuclei (Fig. 5D, left panel). We tested this spatial relationship by placing a counting grid of concentric circles (radius increased in 2 μm increments) over the center of mass for normal pyknotic bodies in H9 and Aβ42 cultures. The area of 7A1a staining in each ring relative to the distance from the center of mass is plotted as a histogram in Fig. 5D (right panel). Pyknotic nuclei have more 7A1a staining nearby relative to normal intact nuclei. Surprisingly, this relationship is quite similar for both Aβ42 edited and unedited H9 cultures. This suggests that pyknosis may be caused by aggregated/oligomeric Aβ42 derived from either direct expression or through APP amyloidogenic processing.

#### Synaptic density

A decrement in the number of synapses is a consistent and early AD phenotype that correlates well with cognitive decline, even during preclinical disease stages [40]. Several transgenic mouse models exhibit synaptic deficits, but we are unaware of this phenotype being described in human cell culture models. We stained 34-day old cultures with anti-synapsin 1 antibody (a presynaptic marker) to estimate the number of synapses present in neuronal clusters from the different genotypes. As shown in Fig. 6, all 3 genotypes at this culture stage have a significant number of synapsin positive puncta. There are, however, ~50% fewer synapsin positive puncta in Aβ42 edited samples (p < 0.0147) relative to unedited H9 samples. Aβ40 samples had ~20% fewer synapsin puncta, but did not reach significance. There is thus a graded genotype dependent difference in the number of synapsin puncta at this culture stage: H9> Aβ40>> Aβ42. We did not distinguish if the Aβ42 synaptic deficiency was due to decreased synaptogenesis or increased synaptic loss. Our results establish that synaptic number is reduced to a greater extent in Aβ42 compared to Aβ40 cultures a result that is consistent with the concept that Aβ negatively affects synaptic capacity [40].

**Fig. 6.**
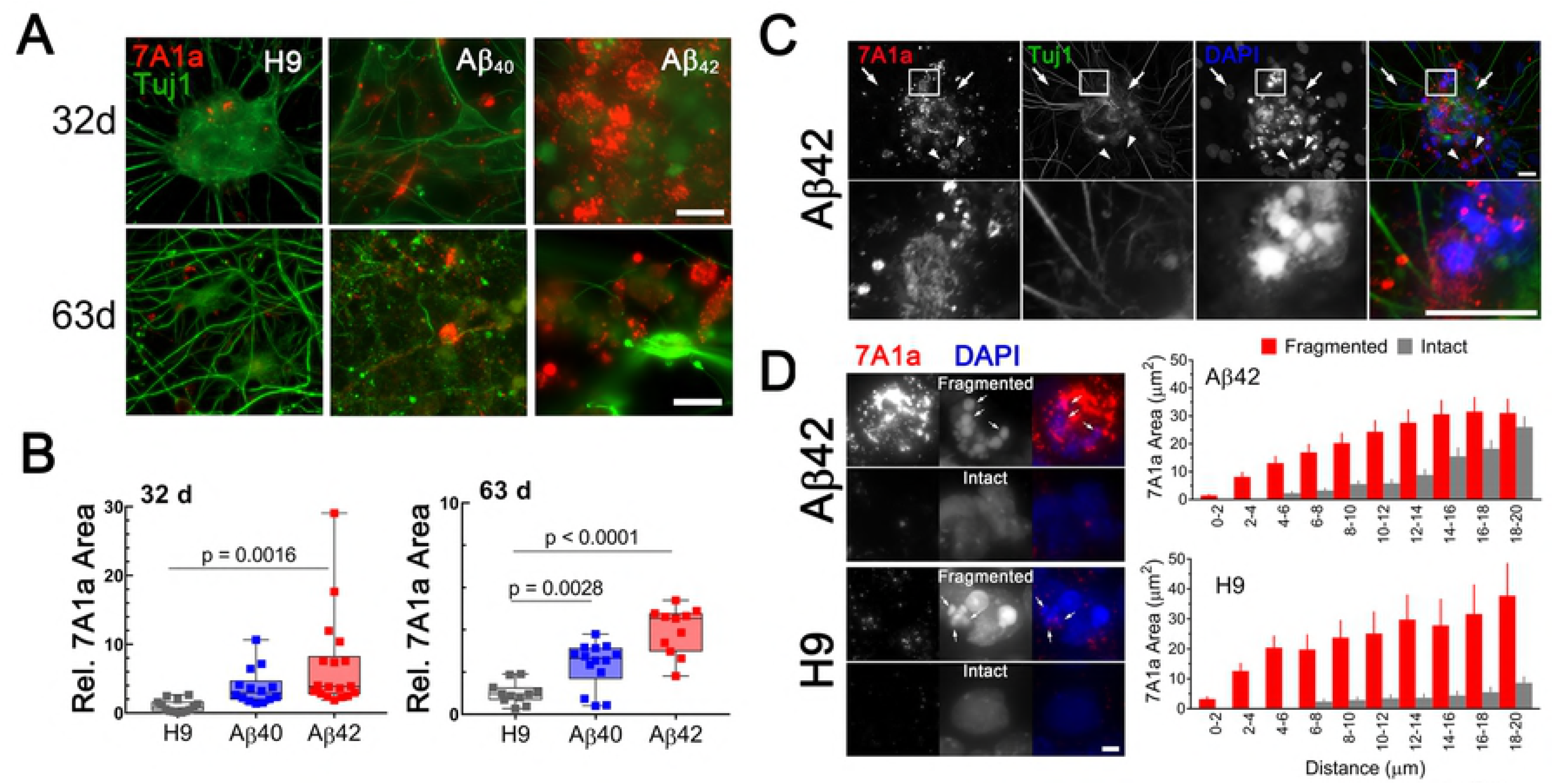
Aβ42 edited NCs have fewer synapsin1 stained puncta in 34-day old cultures. Images are maximum intensity projections of 3 adjacent 0.05 μm spaced optical sections stained with anti-synapsin1 (synaptic marker, green) and anti-NeuN (mature neurons, red) antibodies and DAPI (total cells, blue). Synapsin1 positive puncta were counted in individual NCs from 3 different differentiations normalized DAPI and analyzed (ANOVA, Dunnett corrected). The number of synapsin1 puncta was significantly less for Aβ42 cultures. The relative number of puncta ±SEM were: H9 = 1±0.198, Aβ40 = 0.618±0.065, Aβ42 = 0.492±0.081. Data was from 3 independent differentiations. Scale bar = 20 μm.

#### Progressive ND

AD is a chronic progressive disease with end stage neuronal cell death, a phenotype that has been particularly difficult to document in most current experimental models. We used a fluorescent live/dead assay to assess neuronal viability at 3 different culture ages. Representative morphological and fluorescent images of the same field are shown in Fig. 7 (top). Despite a normal morphologic appearance and similar numbers of neurons in 10-day old cultures, we found a slightly higher proportion of ethidium homodimer fluorescence (dead cells) in Aβ42 cultures even at this early culture stage (Fig. 7, bottom). At an intermediate culture age (34-39 days) when Aβ42 neuronal clusters have significantly fewer synapsin puncta, the relative ethidium homodimer fluorescence was greater in Aβ42 compared to either Aβ40 or H9 cultures. When maintained for longer times (i.e. > ~60 days) both Aβ40 and Aβ42 edited cultures exhibit significantly more relative ethidium homodimer fluorescence compared to unedited H9 cultures. Since most cells under our culture conditions are neurons (~70-90% Tuj1 positive), we conclude that editing results in progressive ND. This phenotype appears at a faster rate for Aβ42 cells relative to Aβ40 cells and is dependent on editing. No viable cells remained in edited culture older than 120 days while H9 cultures still appeared healthy even after 266 days. This edit specific progressive ND also appears to be chronic because of the extended time necessary for its elaboration.

**Fig. 7.**
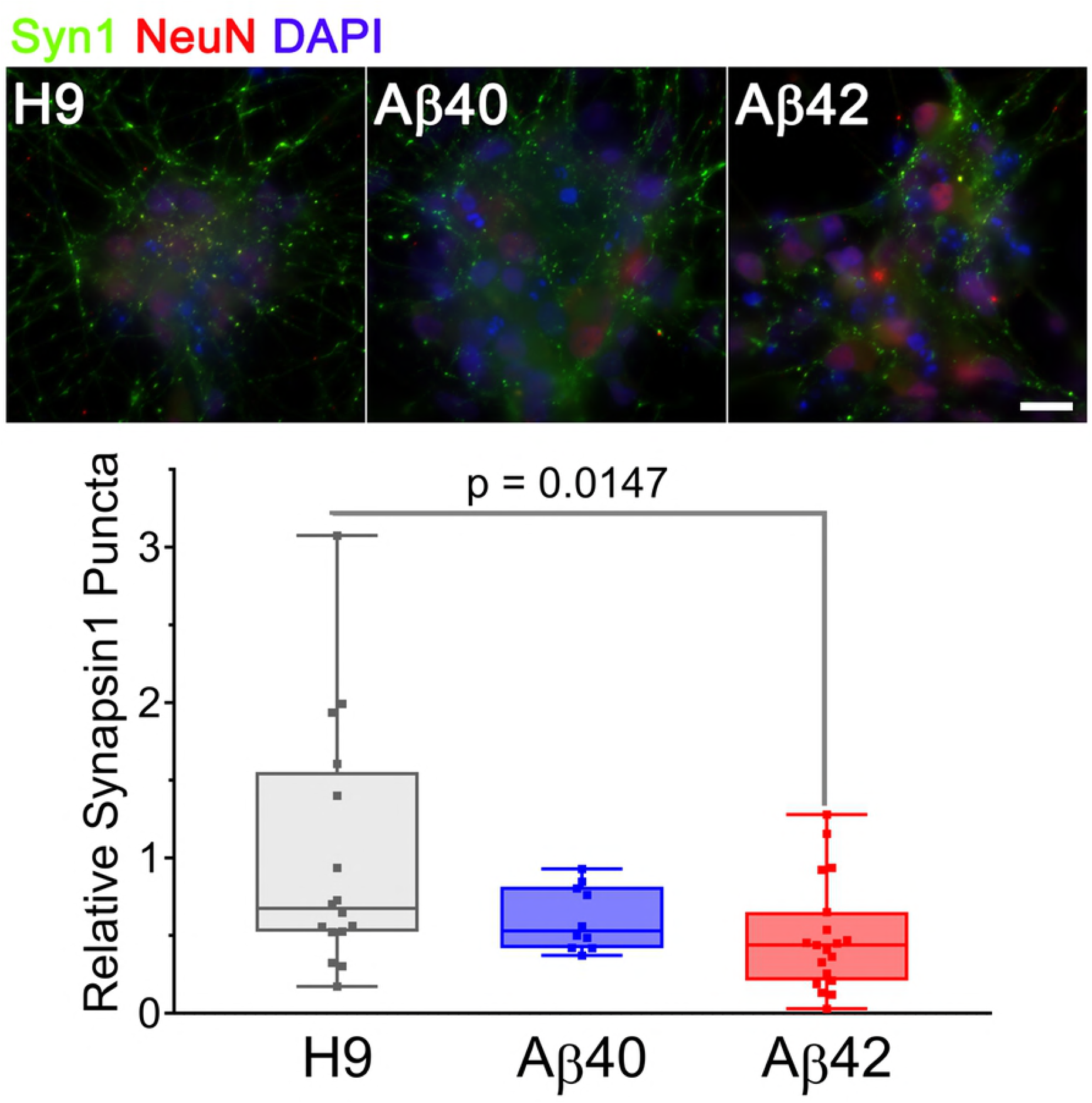
Aβ42 and Aβ40 edited cultures undergo progressive ND. (**Top**) Hoffman extended depth-of-field images (left) with a corresponding fluorescent maximum intensity projection (right) at 3 different culture ages. Green fluorescence (calcein-AM) and red fluorescence (ethidium homodimer was used to estimate live or dead cells. (**Bottom**) Quantitation of relative ratio of dead/live cells. Relative to unedited H9 cultures there are significantly more dead neurons in Aβ42 samples at all three culture ages (ANOVA, Dunnett corrected). Aβ40 samples have significantly more dead neurons but only in cultures older than 60 days. Mean values (±SEM) for 10-day old samples were: H9 = 1.173±0.289, Aβ40 = 3.4±0.643, Aβ42 = 4.49±1.471; for 34-39-day old samples: H9 = 23.9±3.226, Aβ40 = 20.79±2.025, Aβ42 =35.00±2.974 and for >60-day old samples: H9 = 16.02±1.612, Aβ40 = 32.36±3.016, Aβ42 = 38.46±1.588. Each data point represents an individual NC collected from a total of 8 individual differentiations. The line inside the box is the median and the whiskers are the range. Scale bar= 100 μm.

#### Endolysosomal pathway phenotypes

Dysfunction of the endolysosomal pathway, plays an important role in several neurodegenerative diseases, including AD [41]. Pathway dysfunction is a consistent feature of several animal and cellular AD models [23,42] as well as an early phenotype in AD [43] and can be inferred by accumulation of an abnormal number or size of characteristic vesicles.

Using vesicle type specific antibody staining we counted the relative number of punctate vesicular structures in neurons. Fig. 8 presents representative images and analysis for 38 and 62-day old cultures stained with anti-lysosomal associated membrane protein 1 (LAMP1) antibody. There was a ~2-fold increase in LAMP1 positive puncta in 38-day old Aβ42 cultures relative to either Aβ40 or unedited H9 cultures. This finding agrees with the reduced neuronal viability in Aβ42 samples and synapsin1 puncta at this culture stage. In older cultures (62 days) the number of Lamp1 puncta relative to unedited H9 cells was decreased ~60% in Aβ42 and ~50% in Aβ40 samples (although not significant). This decrease thus correlates with ND present in both edited genotypes in older cultures. Abnormal accumulation of lysosomal related vesicles may thus be a consequence of direct Aβ expression in human neurons.

**Fig. 8.**
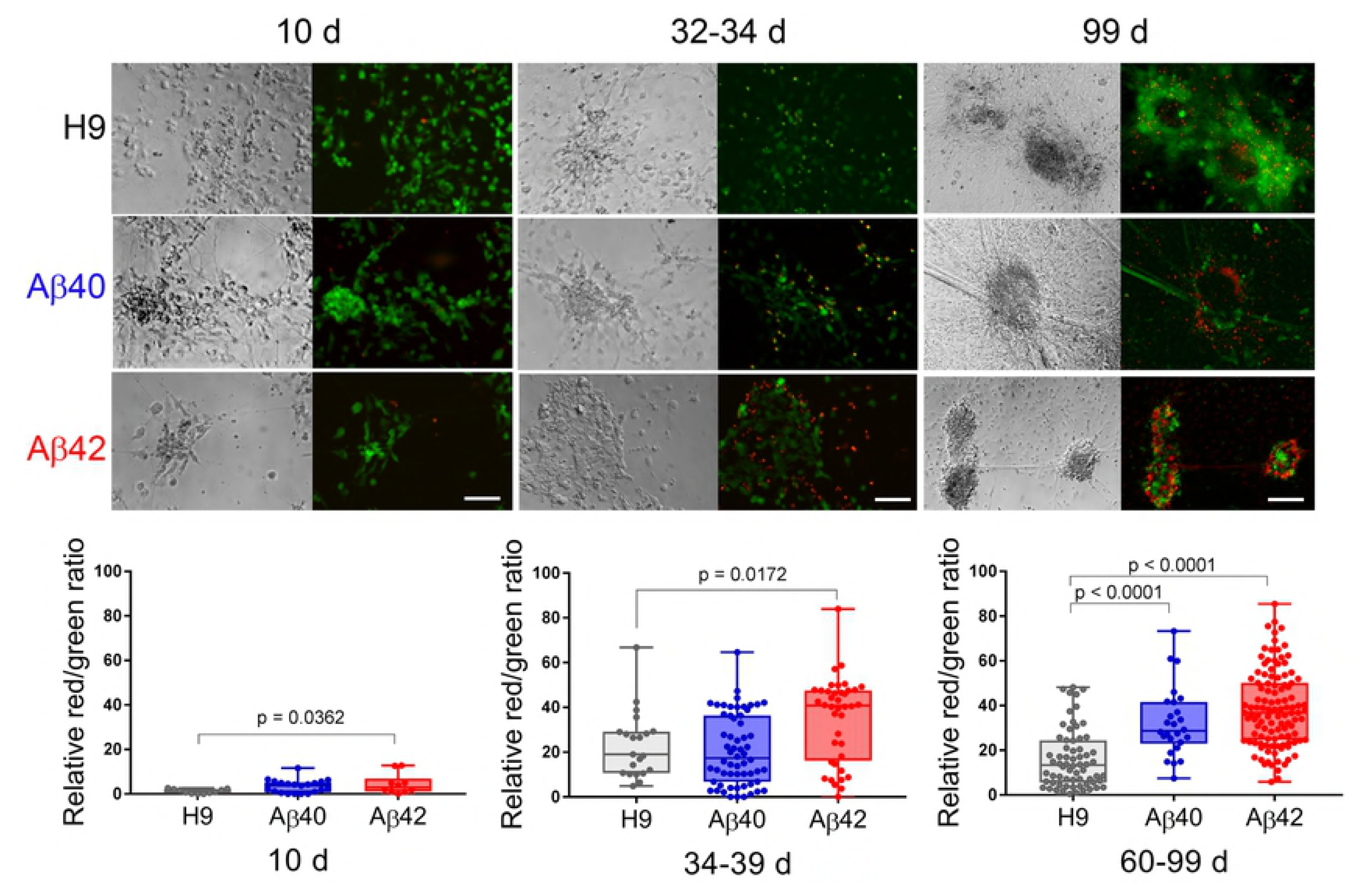
The number of LAMP1 positive vesicles is affected by editing. (Left), fluorescence maximum intensity projections of 2 adjacent optical sections stained with anti-LAMP1 antibody (green) or DAPI (blue) at 2 different culture times. (Right) The relative number of LAMP1 positive puncta (normalized to DAPI) in individual NCs was greater in 38 day old Aβ42 samples relative to H9. In 62 day cultures both Aβ42 and Aβ40 (not significant) samples have fewer LAMP1 objects relative to H9 (ANOVA, Dunnett corrected). Data are from 3 independent differentiations of 38 day cultures and 2 independent differentiations of 62 day cultures. Mean values (±SEM) for 38 day samples were: H9 = 1±0.139, Aβ40 = 0.84±0.059, Aβ42 = 2.064±0.142 and for 62 day samples H9 = 1±0.205, Aβ40 = 0.553±0.106, Aβ42 = 0.453±0.101. The line inside the box is the median and the whiskers are the range. Scale bar = 10 μm.

The number of Rab5 stained puncta, a marker for early endosomes necessary for vesicular maturation leading to lysosomal fusion [44] is shown in Fig. 9A. The pattern is similar to LAMP1 puncta. There was a significant increase in Rab5 puncta in 38-42 day old Aβ42 relative to H9 samples and a non-significant increase in Aβ40 samples. Both edited genotypes also showed a significant decrease in Rab5 puncta in older 63-day cultures. Fig. 9B additionally shows puncta counts for Rab3A, a synaptic vesicular gene important for regulating normal synaptic neurotransmission [45] and LC3B, an autophagosome vesicle marker necessary for delivering mature autophagic/endosomal vesicles to lysosomes for cargo digestion which has been associated with AD [46]. The number of Rab3A puncta were not significantly different among any of the genotypes in 43-day old cultures but both genotypes exhibit a reduction in 63 day old cultures. Both Aβ40 and Aβ42 samples had a reduction in LC3B puncta in 43 day old cultures (only Aβ40 was significant) as well as in 63 day cultures.

**Fig. 9.**
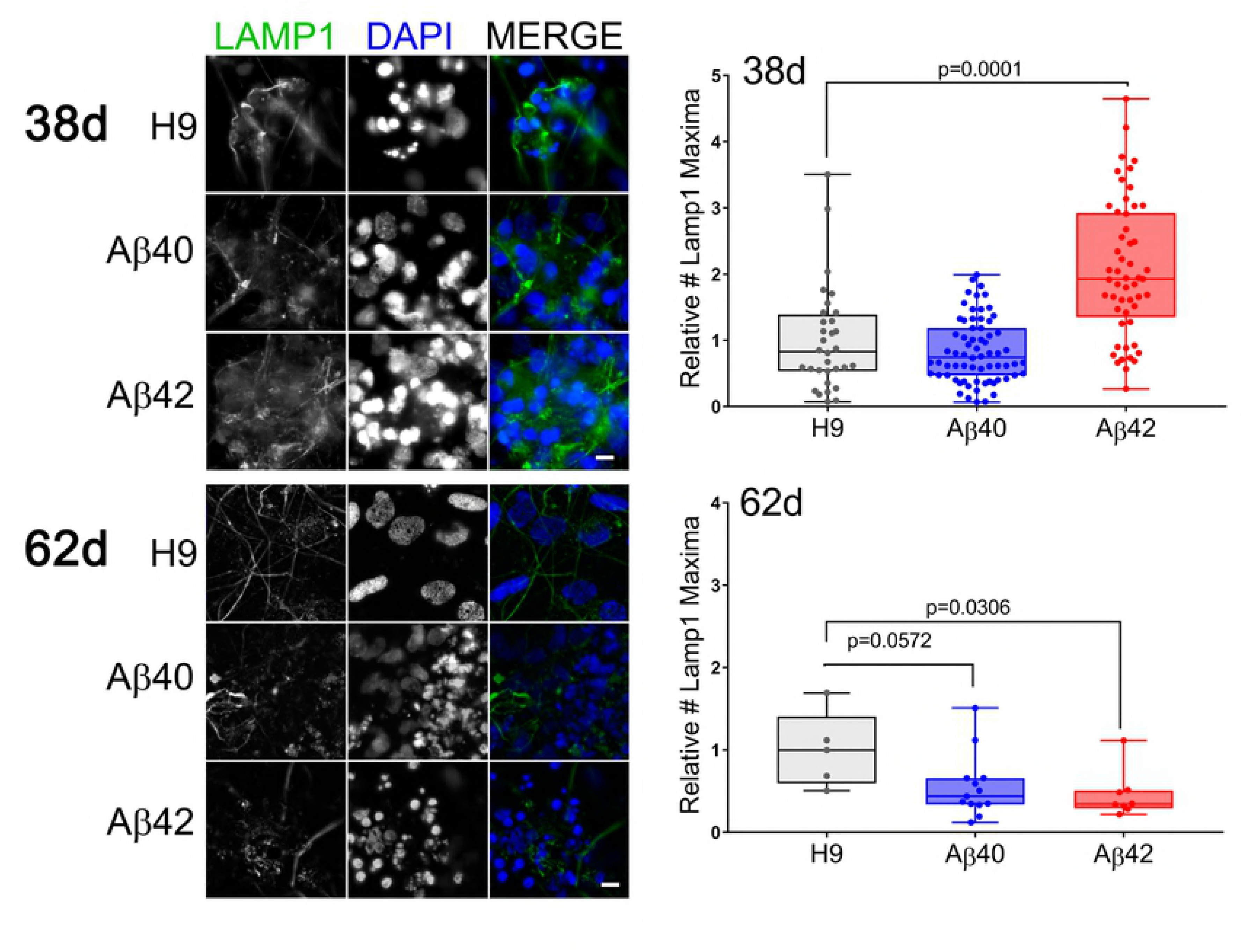
The number of other endolysosomal vesicles is affected by editing. Rab5 positive objects in NCs is greater in 38-42 day old Aβ42 samples. At a later culture age (62 days) both Aβ40 and Aβ42 samples have fewer LAMP1 objects. **A**. (**Left**), maximum intensity projections of 2 adjacent optical sections (1 μm spacing) stained with anti-Rab5 antibody (early endosome marker, red) and DAPI (blue). (**Right**) The relative number of Rab5 puncta (normalized to DAPI) is greater in 38-42 day cultures for Aβ42 edited samples and less for both Aβ42 and Aβ40 edited samples in 63 day cultures (ANOVA, Dunnett corrected). Data is from individual NCs from 3 independent differentiations for 38-42 day cultures and 2 independent differentiations for 63 day cultures. Mean values (±SEM) for 38-42 day samples are: H9 = 1±0.1448, Aβ40 = 2.39±0.2767, Aβ42 = 4.80±1.333 and for 63 day samples are: H9 = 1±0.1584, Aβ40 = 0.586±0.071, Aβ42 = 0.4198±0.0341. **B**. The relative number of Rab3A and LC3B puncta were more variable but both decreased primarily in older cultures. Individual NCs from 2 independent differentiations were stained with either anti-Rab3A (synaptic vesicle associated marker) or LC3B (autophagosome marker) antibody. There was a decrease in LC3B objects in Aβ40 samples at 43 days and a decrease in both Aβ40 and Aβ42, as well as LC3B puncta, in 63 day cultures (ANOVA, Dunnett corrected). Bars are mean ±SEM, N=5-20). Scale bar = 20 μm.

Taken together these results indicate that endolysosomal pathway dysfunction is associated with Aβ edited samples and that Aβ42 samples appear to be affected at earlier times and to a greater extent than Aβ40 samples. These changes are not likely due to changes in gene expression for key vesicular genes since qRT-PCR analysis did not find any genotype specific changes in gene expression (see Supplemental Fig. S2). Since we are directly expressing Aβ in edited cultures, these potential AD related phenotypes are also likely to be largely independent of APP amyloidogenic processing which occurs in large part within endolysosomal vesicles [47].

#### Somal accumulation of phospho-tau

Accumulation of hyperphosphorylated tau (p-tau) and formation of paired helical filaments is a pathological hallmark of late stage AD [7]. We examined the immunocytochemical staining of 62 day old cultures with an antibody specific for tau phosphorylation on serine 244, known to be increased by Aβ [48]. Image analysis of the total area of p-tau staining (normalized to DAPI) was not significantly different between Aβ42 and H9 cultures. The cellular distribution of the staining, while consistent with a redistribution of p-tau from neural processes to cell soma (Fig. 10, top) was only observed in late stage Aβ42 cultures already exhibiting significant ND. The apparent “redistribution” may thus be due to an accompanying decrease in neural processes of dead or dying neurons. The level of tau expression (MAPT gene product) measured by RNA-Seq analysis was relatively similar among the genotypes in earlier age cultures (Fig. 10, bottom). Increased hyperphosphorylation and redistribution of tau which has previously been observed in iPS AD cell models [25,26] but does not appear to be a significant phenotype of direct Aβ expression.

**Fig. 10.**
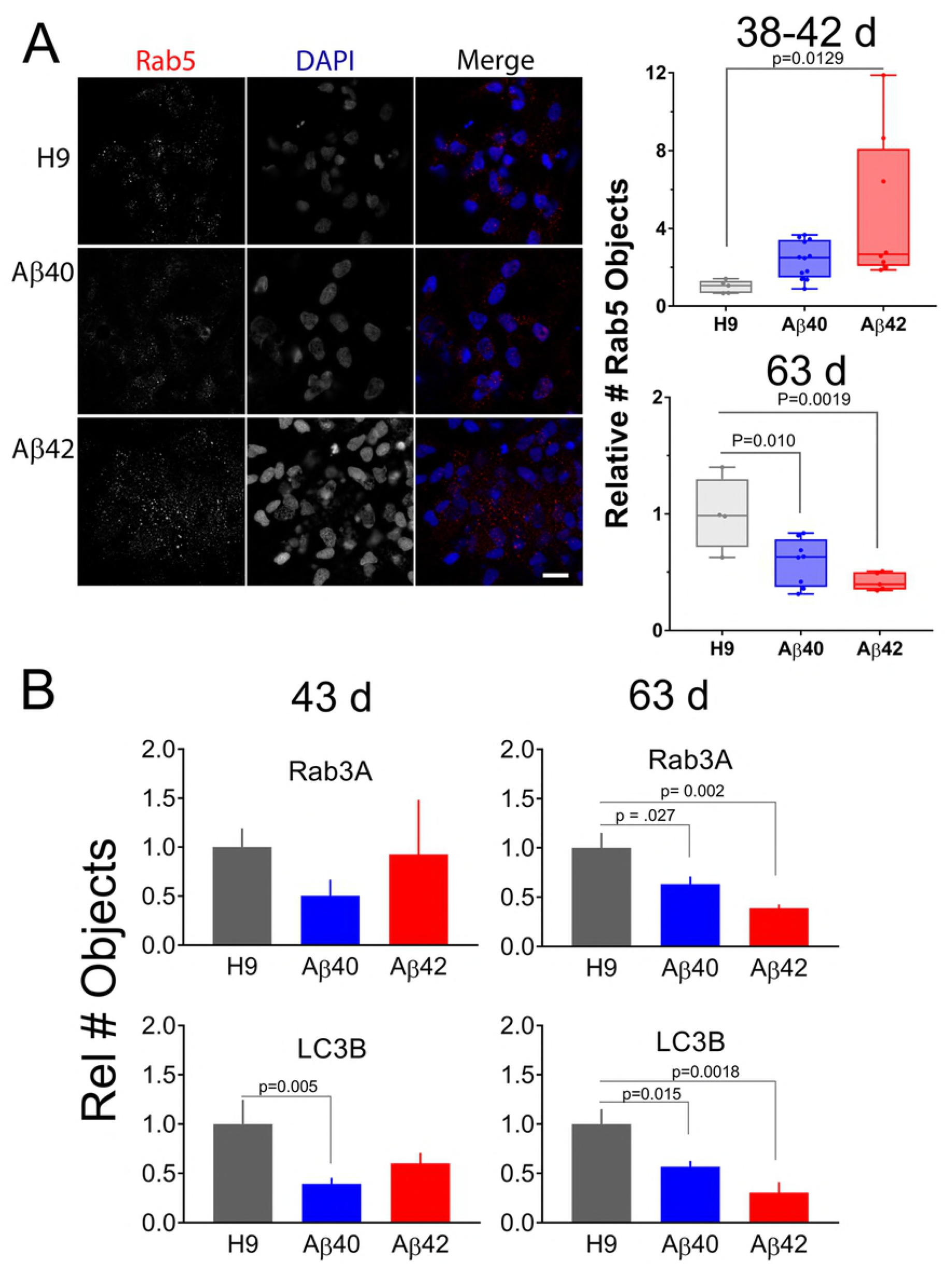
Older Aβ42 cultures show apparent distribution of phospho-tau in cell soma compared to H9 cultures where it is localized in neurites. (**Top**) Fluorescence images from 3 representative fields for each genotype taken from a 62 day old culture stained with anti-phospho-tau antibody (green) and DAPI (blue). This apparent difference is likely due to a significant decrease in neurites on dead or dying cells present in Aβ42 cultures rather than a redistribution of signal. The area of phospho-tau staining (normalized to DAPI) was not significantly different between H9 and Aβ42 samples (p=0.9078, N>15, t test). Scale bar = 20 μm. (**Bottom**), RNA-Seq analysis indicates no significant difference in relative MAPT expression (coding for tau) among the genotypes (ANOVA, Dunnett corrected). Data points are from independent RNA-Seq samples (±SEM).

### Aβ-dependent differential gene expression

The edited cell lines present a particularly favorable opportunity for whole transcriptome RNA-Seq analysis to identify differentially expressed genes (DEGs) that may be mechanistically linked to Aβ-dependent ND. They are not confounded by uncontrolled amyloidogenic APP proteolysis, overexpression of non-Aβ fragments and are near isogenic. We performed RNA-Seq expression using mRNA isolated from 36-38-day old cultures. This is a stage where phenotypes are either exclusive (i.e. reduced number of synapses, reduced neuronal viability and increased accumulation of lysosomes and endosomes) or more penetrant (greater accumulation of aggregated Aβ) for the Aβ42 editing compared to Aβ40 editing. RNA isolated from 3 independent H9 culture samples served as the reference control to identify DEGs for each edited genotype. All three genotypes are heterozygous for the major sporadic AD risk allele (i.e. ε4/ε3) and thus in an appropriate human genetic context relevant to a large proportion of SAD cases [49].

We tested differential expression for 18,259 genes (i.e. genes that had an FPKM > 0.1 in 50% of samples). Results of hierarchical clustering along with an expression heat-map for the batch centered sample medians of individual samples are shown in Fig. 11A. The 4 Aβ42 samples cluster together on the same branch of the dendrogram. One Aβ40 sample (#31.1) clusters adjacent to the Aβ42 group while the other (#41.1) appears more like unedited H9 samples indicating that whole transcriptome expression is more similar among individual Aβ42 edited samples relative to either Aβ40 or unedited H9 samples which agrees with phenotypic penetrance at this culture age. DEGs may thus be mechanistically associated with Aβ42-dependent affected pathways related to these phenotypes. We defined DEGs by first identifying genes that vary between Aβ42 vs H9 and then filtering genes with a similar directional change for Aβ40 that using a more liberal criteria (to avoid keeping genes marginally not significant in the Aβ40 vs H9 comparison) (see **Supplemental Methods and Data** for full details). All 93 DEGs for the Aβ42 vs H9 comparison are shown in Fig. 11B as a fold change (FC) heat map. There were 23 UP and 70 DN (down) regulated genes which were used for functional/annotation enrichment analysis. This number is rather small compared to numerous other AD related studies of DEGs in patient samples or even iPS cell lines where thousands of DEGs are often identified [23,50–52]. Note that the directional FC (fold-change) was similar for most genes in the Aβ40 samples compared to Aβ42. The Pearson correlation coefficients for log2 ratios of all Aβ42 vs H9 compared to Aβ40 vs H9 genes was 0.5434 (all genes, linear-regression p-value < 0.0001, Fig. 11C, top), and the correlation coefficient for the DEG FC values is 0.3183 (differentially expressed genes, linear regression p-value = 0.0019, Fig.11C, bottom). Aβ dependent changes in gene expression thus appear similar for Aβ42 and Aβ40 samples. A complete list of all detected genes, the genotype specific average log2 RPKM values, log2 ratios of the H9 DEG comparisons, FC values, statistics and DEG status is included as **Supplemental Table S1**.

**Fig. 11.**
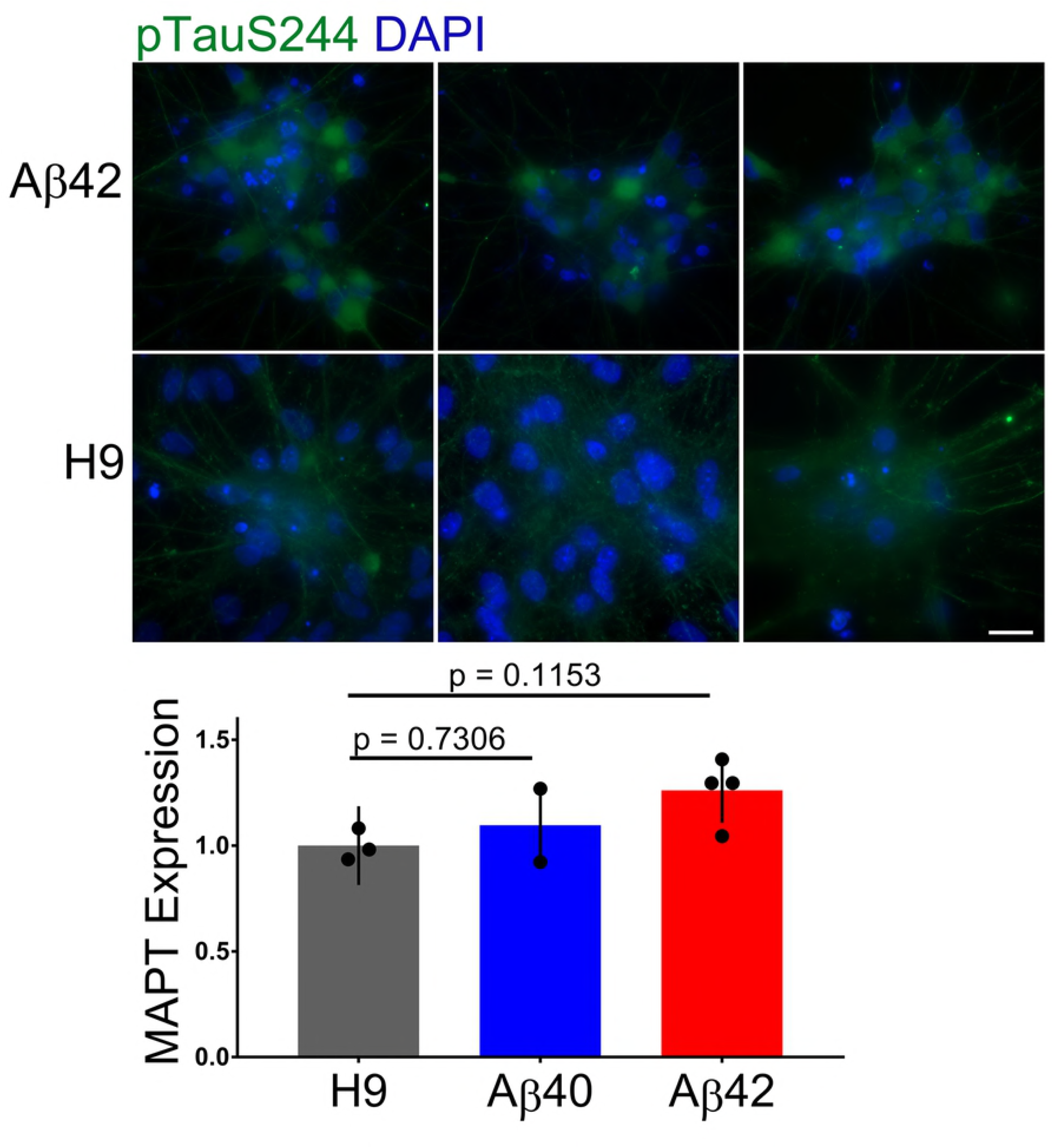
Differentially expressed genes in 34-day old cultures. **A**. Cluster analysis of differentially expressed genes (Pearson dissimilarity metric). Aβ42 samples cluster together while Aβ40 and H9 samples overlap. **B**. Heat map of significant DEGs from RNA-Seq analysis of Aβ42 vs H9 comparison and Aβ40 vs H9 comparison. Up (UP) regulated genes (red) and down (DN) regulated genes (green) were sorted by the magnitude of the indicated fold change (FC) values for the Aβ42 vs H9 comparison. There is a general correspondence in the directional FC values with a relatively larger FC in the Aβ42 samples. **C**. Pearson correlation confirms significant co-variation of the log2 ratios of all genes (top, significant DEGS in color) as well as the FC values for significant UP and DN regulated genes (bottom).

GO enrichment analysis of the UP and DN regulated genes for the Aβ42 vs H9 comparison did not identify functional enrichment for UP genes after correcting for FDR. The statistical power of this approach, however, is likely limited when using a small number (23) of input DEGs. For DN genes, however, 13 out of 70 (19%) were related to cilia functions and were significantly overrepresented (i.e. FDR<0.05) (CCDC114, CFAP100, CFAP126, CFAP45, CFAP70, DAW1, DNAAF1, DNAH11, DNAI2, SPAG17, STOML3, TEKT1, USH2A). Interestingly, 5 of these “cilia” genes (DAW1 DNAH11 DNAI2 GDA TEKT1) were also differentially expressed in a hippocampal AD vs non-AD RNA-Seq study [50] suggesting that cilia related pathways may also be affected in AD. Using unadjusted p-values, microtubule and cytoskeletal genes were also over represented (CCDC114, CFAP100, CFAP126, CLIC5, DNAAF1, DNAH11, DNAI2, GAS2L2, PARVG, SPAG17, TEKT1, USH2A) as well as genes associated with vesicle lumen (COL11A1, COL8A1, ERP27). Overrepresented molecular functions included neurotrophin receptor associated terms (NTRK1) and peptidase regulatory roles (CD109, SERPINA3, SERPIND1). The complete GO results are included in **Supplemental Table 2**.

We also used GATHER (http://changlab.uth.tmc.edu/gather/gather.py) to broaden the search for relationships/pathways in the Aβ42 DEGs. Two GO terms were statistically significant for UP genes (FDR<0.05): *GO:0007267*: cell-cell signaling (ADRA1B, CPNE6, CXCL14, MME, TNFSF10, UTS2) and *GO:0007154*: cell communication (ADRA1B, COL19A1, CPNE6, CXCL14, DKK1, GRP, HAPLN1, MME, STAC2, TNFSF10, UTS2). DN genes included two overlapping GO terms: GO:0015698: inorganic anion transport and GO:0006820: anion transport (CLIC5 COL11A1 COL8A1 SLC12A1). KEGG pathways with an FDR<0.25 included hsa04080: Neuroactive ligand-receptor interaction (ADRA1B, GRP, UTS2) and hsa05010: Alzheimer’s disease [MME] for UP genes. DN genes were hsa04512: ECM-receptor interaction (COK11A1, FNDC1). Complete GATHER results are included in **Supplemental Table S3**.

GSEA KEGG analysis (http://software.broadinstitute.org/gsea/index.jsp) is an additional way to discover potential pathway relationships and are not limited by by using a small list of input genes since input can be a rank order list of FC values for all detected genes. We performed GSEA using a rank ordered FC list (18,233 genes) and compared these to all KEGG pathways. The Aβ42 vs H9 list identified 118/170 KEGG gene sets that were upregulated. Twenty-two had a nominal p value <0.05 and 3 of these had an FDR <25%. The top scoring KEGG pathway was NEUROACTIVE LIGAND RECEPTOR INTERACTION (hsa04080, Normalized Enrichment Score=2.12, p<0.01, FDR=0.014). Our gene list included 79 of the 219 (36%) genes in this pathway suggesting widespread changes in neuroactive ligand receptor signaling was a consequence of direct Aβ42 expression. This can plausibly be related to the DN regulated expression of “cilia” related genes since primary cilia in neurons are believed to be a major organelle signaling hub known to express a host of neuroactive ligand receptors [53]. No KEGG pathways reached significance (FDR<0.05) for DN genes or for a separate analysis of ranked Aβ40 vs H9 DEG FC values. Summary results for the top 20 GSEA KEGG pathways for Aβ42 vs H9 genes along with details of the KEGG NEUROACTIVE LIGAND RECEPTOR INTERACTION pathway are included in **Supplemental Table S4**.

We also analyzed DEGs using Ingenuity Pathway Analysis (IPA). Remarkably, for the Aβ42 vs H9 comparison, the highest and lowest z scores were obtained for the functions “Increased Neuronal Cell Death” (z = 1.658) and “Decreased Memory” (z = −2.213), two biological processes with obvious relevance to AD. Fig. 12 shows the individual DEGs identified by this analysis color coded by intensity for FC values. The “Decreased memory” and “Increased neuronal cell death” pathways are connected through the overlap of DKK1 and NTRK1. An IPA analysis for disease related pathways returned the “Neuroprotective Role of THOP1 in AD” as the top scoring canonical pathway (p = 9.95 E-3). This pathway was also significant for a hippocampal DEG analysis of LOAD RNA-Seq data [50]. Thimet oligopeptidase (product of THOP1) is reportedly neuroprotective for Aβ toxicity in cortical neurons and can degrade soluble Aβ but not aggregated Aβ42 [54,55]. The DEGs in the Aβ42 vs H9 comparison represent only a small fraction of the 40 genes in this pathway. They were MME (aka NEP, neprylisin) and SERPINA3 (aka ACT) (indicated on the bottom of Fig. 12). MME is not directly related to decreased memory in IPA, but is included because of its potential indirect relationship through GRP [56]. SERPINA3 is a member gene of the “Neuronal cell death” category in IPA and both genes are part of the extracellular arm of the THOP1 in AD pathway in IPA. MME, is an Aβ degrading enzyme with increased expression in the Aβ42 edited cells and SERPINA3, is a serine protease inhibitor with decreased expression which co-localizes with Aβ in AD plaques [57].

**Fig. 12.**
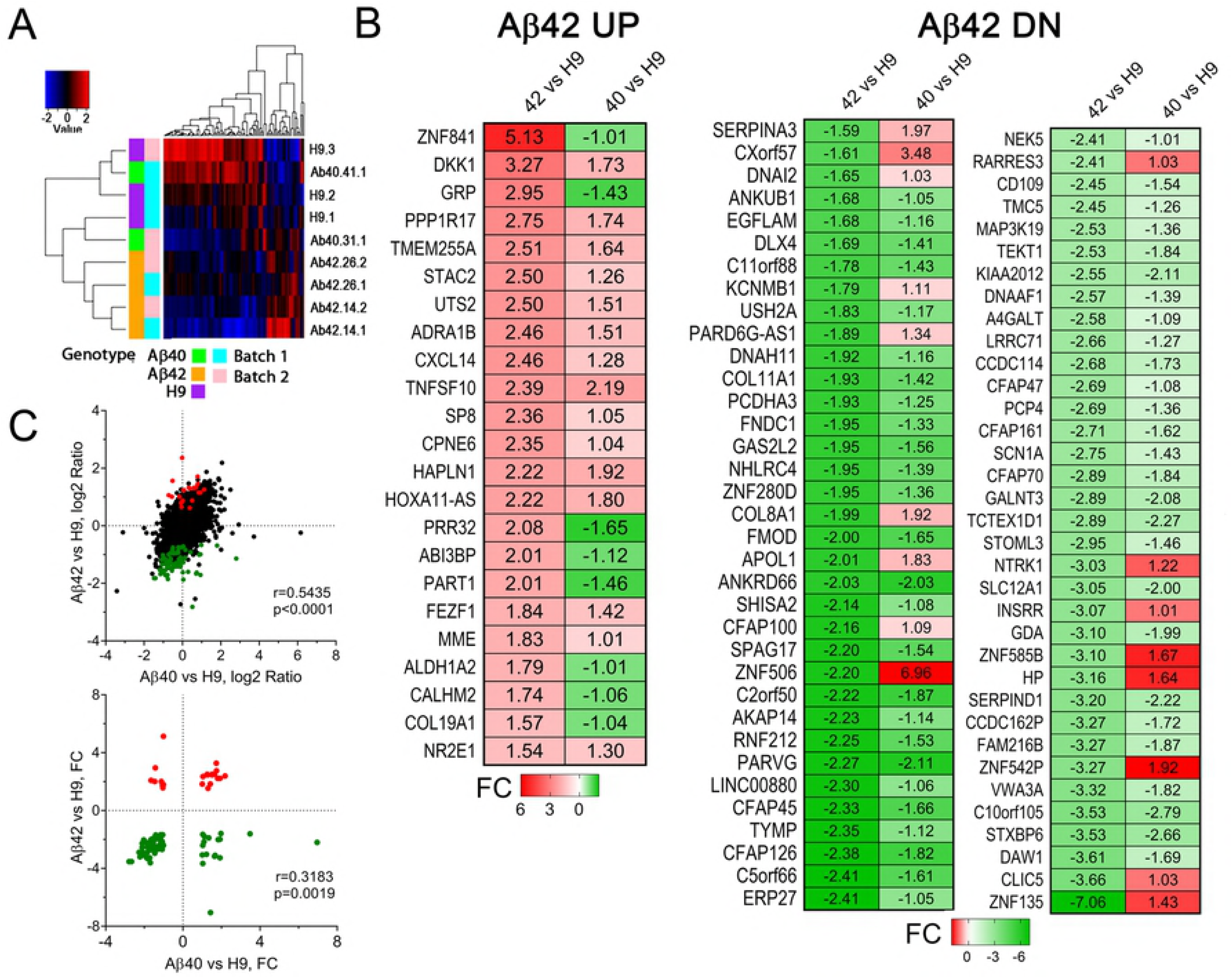
DEGs are potentially related to Alzheimer’s relevant pathways and functions. (**Top**), IPA pathway analysis of DEGs in the Aβ42 vs H9 comparison identified “decreased memory” (z = −2.213) and “increased neuronal cell death” (z = 1.658) as the lowest and highest scoring functional pathways. Individual genes are shown as graphic symbols representing molecule type and color coded by FC values (red=UP, green=DN). (**Bottom**), the most relevant IPA disease related canonical pathway was “Neuroprotective role of THOP1 in Alzheimer’s disease” (p = 9.95 E-03, overlap = 2 of 40 total genes in this pathway). The pathway genes were MME (aka NEP, neprilysin) an Aβ degrading metalloproteinase and SERPINA3 (aka ACT, alpha-1 antitrypsin) a protease inhibitor found in AD plaques (Top, circled in blue). MME is not directly included in the IPA “decreased memory” function but can potentially be indirectly related through its relationship to GRP.

There is no general consensus regarding a “signature” set of AD related DEGs, especially those that related to early LOAD pathogenic mechanisms making it challenging to relate our expression data with patient samples likely to contain signals from many different non-neuronal cell types, co-morbidities and many complex combinations of genetic variance. Nevertheless we did find some encouraging comparisons. For example, the GeneCards database (https://www.genecards.org) has 6,672 genes identified as “Alzheimer’s related genes”. This is a rather large list not restricted to DEG analysis but also including GWAS hits as well as other types of associations. For our UP genes we found 10/23 (43%) that overlapped (TNFSF10, DKK1, GRP, CALHM2, MME, ALDH1A2, CXCL14, PPP1R17, TMEM255A, HAPLN1) and 17/70 (24%) DN genes (SHISA2, DNAH11, SERPIND1, SCN1A, APOL1, HP, ERP27, SERPINA3, STXBP6, CFAP70, PARVG, GDA, PCP4, NTRK1, TMC5, STOML3, RARRES3) suggesting some potential AD relevance. A recent RNA-Seq analysis of hippocampal tissue from SAD vs non-SAD patient samples [50] identified 2,064 DEGs. We found only 3 out 23 (13%) UP genes overlapped (not statistically significant; Fisher’s exact test, p=0.46) (HAPLN1, CPNE6, TNFSF10). In contrast, 22 out of 70 (31%) DN genes overlapped (Fisher’s exact test, p = 2.5×10^−6^) (DAW1, FAM216B, GDA, TCTEX1D1, PCP4, CCDC114, LRRC71, A4GALT, MAP3K19, TEKT1, CD109, TMC5, RARRES3, LINC00880, PARVG, ANKRD66, FNDC1, DNAH11, C11orf88, ANKUB1, DNAI2, SERPINA3). Four of these DN genes had an opposite directional FC, while others agreed with our DEGs. This significant overlap suggests that DEGs in our Aβ-dependent neuronal model may thus have relevance to the AD, including the possible involvement of cilia dysfunction as mentioned above.

## Discussion

The plethora of genes, molecules, cell types and pathways implicated in extensive AD patient and experimental model organism studies have not yet identified critical factors that initiate and sustain the progressive clinical and pathological decline characteristic of this neurodegenerative disorder. Many investigators believe that Aβ accumulation plays a key role in initiating pathogenic processes, however the specific aggregated/oligomeric state responsible is controversial and we lack a clear understanding of mechanisms and pathways that link Aβ to ND. Reasons for this are twofold: AD is an extremely complex disorder and most experimental models do not exhibit progressive ND as a phenotype [4]. Interesting exceptions to this phenotypic deficiency are mouse [11–14] or invertebrate [10] models that directly overexpress Aβ rather than relying on its production via APP amyloidogenic proteolysis. Aβ42 direct expression models all exhibit chronic progressive ND. Human iPS AD models appear to be a promising way to experimentally investigate AD mechanisms in a human genetic context [22,58–60] but unfortunately they also fail to progress to ND.

In the current study, we used genomic editing to obtain isogenic hES cell lines that differ only in a single allele of the normal APP locus. This approach permitted a comparative analysis of phenotypes relative to unedited parental near isogenic cells, as well as independent analysis of direct expression of an equivalent amount of Aβ40 or Aβ42 since both edited alleles are under control of the endogenous rather strong APP promoter. We thus expected the strength and timing of direct expression would match APP itself but this was not the case. Developmental timing of expression agreed well with APP however expression levels of edited alleles were both ~30 fold lower. Our model is thus significantly different from other direct expression models where strong exogenous promoters are used that may complicate phenotypic interpretation because of potential overexpression artifacts. Overexpression of FAD genes, or even wild type APP, seems to susceptible to this complication in transgenic mouse AD models [4]. APP proteolysis is a complex process with both amyloidogenic and non-amyloidogenic pathways producing a host of other fragments that exhibit a variety of documented or potential phenotypic consequences [61]. The phenotypic results we describe are due to direct expression and thus not likely to be confounded by non-Aβ peptides. No amyloidogenesis is required for Aβ generation and APP expression is reduced ~50% by our editing strategy.

The near isogenic nature of our cell lines ensures that phenotypic results are additionally not confounded by genetic variance, known to have small but significant cumulative effects on AD risk. This is an uncontrolled variable in some patient derived iPS models. Additionally, our cell lines may have potential relevance to SAD since no AD FAD related mutants [62] were used and the cells all have the APOE ε3/ε4 genotype which is associated with a large fraction of SAD cases [49]. We extensively characterized two independently isolated clones for each edited genotype and did not observe significant phenotypic differences within each edited genotype. It is therefore unlikely that phenotypes are a result of off target effects or low level mutations which may occur during stem cell editing [32].

### Neurodegeneration

One principal finding of this study is that direct expression of secretory Aβ40 or Aβ42 in cultured human neurons is sufficient to result in a host of AD-like phenotypes up to and including progressive ND. Progressive ND has not been adequately modeled in other non-human animal [6] or current human cellular AD models [22,60] and this phenotypic deficiency has led in part to some of the controversy regarding the role of Aβ in AD [20,63]. The amyloid hypothesis [18] proposes that Aβ is a primary driver for a host of downstream pathogenic cascades terminating in ND. Direct Aβ expression is a simple test of this postulate and our results strongly support it, with the caveat that intraneuronal accumulation, rather than extracellular action appears to be responsible for phenotypic changes. Significant experimental evidence supports an intraneuronal site of action for Aβ42 proteotoxicity [64].

Some major competing hypotheses suggest other key non-Aβ mechanisms mediate ND such as tau-hyperphosphorylation, neurofibrillary tangle formation, generation of non-Aβ proteolytic fragments of APP or pathological action of non-neuronal cell types (i.e. astrocytes or microglial). Our results suggest that these AD associated pathologies are not necessary for progressive ND in cultured human neurons. We did not observe neurofibrillary tangles and only a modest tau redistribution in cultures where ND was nearly complete. Additionally, neurons appear to be the near exclusive cell type present in our cultures suggesting that Aβ proteotoxicity is neuronal autonomous. Importantly, direct expression of Aβ is also necessary to generate the phenotypes we describe, including ND, since neurons derived from unedited parental cells could be maintained up to 266 days while no edited neurons survived beyond 120 days. Finally, the ND we observe is chronic and progressive like AD and thus differs from the well-established acute cellular toxicity of non-physiological concentrations Aβ.

AD is not considered a developmental disease, but rather a condition restricted to old age. This presents a significant challenge for stem cell derived culture models. We could detect significant expression of edited Aβ transcripts in stem cell cultures, during the ~1-month long EB process and during the initial stage of neuronal differentiation. However, we did not observe any major morphologic or molecular differences related to this early exposure to Aβ. Stem cell and differentiation markers, neurogenesis and neuronal marker gene expression were similar for all 3 genotypes in 10 day post EB cultures. We did note that cultures of both edited genotypes sometimes appeared to self-organize into neuronal collections at slightly earlier times than unedited H9 cells (i.e. the cells were often closer together and more likely to bear small process at the earliest differentiation stages (i.e. up to ~7-10 days of in neural differentiation media). This is consistent with a reported “neurogenic” effect of ES cells exposed to Aβ [65]. Direct Aβ expression thus does not appear to adversely affect neuronal developmental processes in our culture system or the ability of neurons to self-organize into NCs.

### Phenotypic timeline

The earliest AD-related phenotypic change we observed for edited relative to unedited cultures was a greater accumulation of aggregated/oligomeric Aβ. This was more prominent at earlier times for Aβ42 cultures and correlated with the rate of decreased neuronal viability. Aβ42 samples had small but significant reduction in neuronal viability even at 10 days while Aβ40 samples did not. By 35 days the level of aggregated/oligomeric Aβ was significantly higher in Aβ42 cultures which also had significant more ND. 35-day Aβ40 cultures appear to trend toward reduced viability but did not reach statistical significance. Direct expression of Aβ42 thus is more toxic to human neurons relative to Aβ40. This agrees with the known in vitro and in vivo propensity of Aβ42 to more rapidly form aggregates or oligomers relative to Aβ40 and suggests that accumulation of aggregated/oligomerized Aβ likely explains the differential rate of ND.

Levels of aggregated Aβ42 were maximal in 32-day old Aβ42 cultures, intermediate in Aβ40 cultures and only present in small amounts in unedited cells. Since both peptides are produced under the same genetic control, our data suggest that cellular mechanisms for Aβ removal may operate more efficiently for Aβ40 than for Aβ42 in human neurons, a result consistent with other direct expression models [11,16]. The low levels of aggregated Aβ accumulation in unedited cells do not appear to be progressive and ND is not a prominent feature of these cultures providing additional support for a direct relationship between accumulation of aggregated/oligomerized Aβ and eventual ND.

Interestingly, the aggregated Aβ appears to be primarily intracellular and appears to accumulate initially in small vesicles, a result consistent with early endosomal accumulation of Aβ in AD now believed to be a major site of pathological amyloidogenic processing [64]. Both edited genes contain a normal secretory signal sequence to route peptide production through the normal secretory vesicular pathway, similar to APP. Since we were unable to detect peptide in culture media we don’t know if Aβ peptides were secreted at comparable levels and/or if they were then reinternalized in endocytic vesicles. In other direct expression AD models Aβ42 appears to be preferentially retained or endocytosed by neurons relative to Aβ40 [11,17]. Secretion and reuptake of Aβ has also been suggested in cultured neurons or early stage AD patient samples [66].

With respect to endocytosis, we saw a significant increase in accumulation of both lysosomes (LAMP1 positive structures) and early endosomes (Rab5 positive structures) in Aβ42 edited neurons suggesting abnormalities in these particular vesicular pathways. Early endosomes containing Aβ may potentially mature and fuse with lysosomes which are in part a common intracellular vesicular transport pathway. Altered endo/lysosomal pathway function has previously been reported in early stage AD brain as well numerous mouse AD models and FAD iPS models [36,43]. The low internal pH of maturing endosomes and lysosomal/endosomal fusions would likely promote Aβ aggregation. The 7A1a antibody we used in this study specifically detects aggregated Aβ42 in vesicle compartments in *Drosophila* neurons [17]. The increased number of lysosomes and endosomes was not maintained at longer culture times, possibly because of increased ND at these later stages. This is consistent with proposals for how intracellular Aβ aggregates could eventually form extracellular plaque structures, a defining feature of AD pathology [17,67]. Notably we did not observe extracellular plaque like structures possibly because they may be removed during frequent media changes. A cause-effect relationship between extracellular plaques and AD pathology has not been definitively established.

### Aβ and pyknosis

An interesting correlation we observed was the close spatial association of aggregated/oligomeric Aβ and pyknotic neuronal nuclei. This type of nuclear fragmentation is a defining characteristic of apoptotic cell death, but is also seen in other types of cell death [68]. Notably, this correlation was not strictly dependent on direct Aβ expression since it was observed in unedited samples. Pyknosis may thus be related to low levels of amyloidogenic APP processing in the unedited neurons. The mechanism(s) of neuronal death in AD is not completely established, however, some evidence for apoptosis has been described in more accessible cell culture and animal models [69]. Direct injection of small amounts of Aβ42 peptide or an episome expressing Aβ42 into the cytoplasm results in acute toxicity of primary cultured human neurons. Cell death was mediated through p53/BAX dependent apoptosis and associated with significant evidence of condensed nuclear chromatin [70]. In this same study, Aβ40 was not toxic in contrast to our results. The Aβ40 toxicity we describe, however, only manifests at a significantly later culture time relative to Aβ42 cultures and the cytoplasmic injection results were obtained only a short time after injection perhaps explaining this difference.

### Direct Aβ40 expression suggests a human specific phenotype

The delayed toxicity we observe for Aβ40 diverges significantly from findings in both rodent and *Drosophila* direct Aβ overexpression models where Aβ40 expression appears to be relatively limited in its ability to generate AD-like phenotypes [11,13–15]. In *Drosophila*, high levels of Aβ40 over expression in cholinergic neurons appear beneficial as they extend the lifespan [16]. The absence of Aβ40 toxicity in flies is likely due to increased secretion or intracellular removal relative to Aβ42 which preferentially aggregates within intracellular endolysosomal vesicles [17]. Rodent direct Aβ40 models produce human peptide (from the transgene) in the context of endogenous production of rodent Aβ peptides (from the endogenous mouse gene). These peptides have 3 different specific amino acids that could modify the human peptides ability to form toxic aggregates or oligomers. Different combinations of Aβ peptides are known to interfere with the rate of formation of aggregates/oligomers as well as their structural type [71]. This is consistent with the possibility that shorter Aβ40 peptides (or sequence divergent rodent peptides) could prevent toxicity of Aβ42 in rodent models. The *Drosophila* homologue of APP does not contain any Aβ sequence so it would not be subject to the same process. Additionally, a recent human-mouse hybrid AD model also observed human specific ND in response to mouse neuronal production of FAD derived amyloidogenic Aβ [72].

Our unedited cultures clearly produce small amounts of aggregated Aβ and this process also likely occurs in edited cells (i.e. APP expressed from the unedited allele). It is possible that a small amount of amyloidogenic derived Aβ could act as a “seed” to stimulate additional aggregation/oligomerization of directly expressed Aβ40 and Aβ42. Such a process could account for the ND properties of our model. Further study will be required to examine this possibility. Many AD related studies use incompletely defined oligomers of Aβ isolated from AD brain that seem to have the ability to specifically initiate several AD-like phenotypes [73]. The exact structure of proteotoxic Aβ assemblies are still imprecisely defined, but generally believed to be smaller oligomers. It will be interesting to compare the Aβ structures present in our cultures with those isolated from AD brain.

### Synaptic deficits

We document a deficit in synapsin1 stained puncta in 34-day old cultures which was specific for Aβ42 edited cells at this culture age. Synaptic deficits are an early AD phenotype but have rarely been reported in human AD culture models [22]. Other experimental models attribute synaptic deficits to an increased production of Aβ associated with increased synaptic activity, or a complex relationship to amyloidogenesis (sometimes involving non-Aβ40 or Aβ42 APP derived proteolytic products) or even a tau dependence [40]. The deficit we observe is most likely a direct result of Aβ42 aggregation/oligomerization since amyloidogenic processing is likely to be minimal in edited cultures. Additionally, we do not observe tau related phenotypes until much later culture ages suggesting that the synaptic deficit is not mediated by tau dependent pathogenic processes. Importantly, we did not distinguish between the failure to form synapses or their increased removal which would require additional observations.

### Phenotypic changes appear to be neuron specific

We used H9 ES cells as the parental genotype in this study since they are widely used in many neuronal differentiation protocols, have been successfully edited and have a SAD associated APOε4/APOε3 genotype [49]. An ongoing challenge in constructing relevant neurodegenerative stem cell models is differentiation to specific cell types involved in the disease [60]. Many types of neurons degenerate in AD and we initially tested a differentiation protocol designed to produce an enrichment of cholinergic basal forebrain-like neurons that was used to generate H9 derived neurons susceptible to Aβ42 oligomers [74]. Unfortunately, we could not reliably obtain differentiated neurons that could be maintained in culture for more than a few weeks. Successful generation of cholinergic neurons was also minimal, suggesting that protocols to generate basal fore brain neurons may still need improvement [75].

The work presented in this communication used a protocol originally developed to generate enrichment of limb motor neurons [29], a neuronal type not generally affected in AD. We note however that RNA-Seq data suggests a neuronal population significantly more complex than primarily limb motor neurons. First, low levels of CHAT expression were detected and only ~10-20% of NeuN positive cells were positive for staining with an anti-choline acetyltransferase antibody (data not shown). No muscle cells are present and their absence, especially in longer term cultures would likely have a significant effect on the ultimate state of motor neuron differentiation. We detect expression of GAD1 and GAD2, TH and SLC17A7, suggesting additional non-motor neuron neurotransmitter phenotypes are present. It is likely that we derived a neuronal cell population biased towards caudal, rather than rostral differentiation. For example, caudal HOXB4, HOXB6 and HOXA1 levels were high but not GLRA1 and MNX1. Likewise, some but not all rostral genes agree with a more rostral fate (i.e. high levels of rostral CUX1, SATB2, RELN and DAB1 but low levels of TBR1 FOXG1 and NKX2-1). A direct comparison of these two general classes of neurons in iPS cultures established greater early stage AD-related phenotypic elaboration in rostral neurons, however the differences were not large and were possibly related to differences in the rate of amyloidogenesis rather than the direct expression we use here [76].

It is clear from RNA-Seq data that our cultures are primarily neuronal (ie. high level expression of DCX, TUBB3 and MAPT) in good agreement with ~90% of cells reliably staining with antibody to either NeuN, Tuj1 or DCX. We also detected very low and marginally significant expression of astrocytic markers ALDH1L2, GFAP; oligodendrocyte marker OLIGO2 and microglial marker TREM2 and AIF1 but did not observe any cells with characteristic morphology of these glial cell types. One necessary modification we made to the original differentiation protocol [29] was a weekly 24 hour exposure to EdU to suppress mitotic cell overgrowth (continued up to ~50-60 days of differentiation). This treatment is likely to eliminate the bulk of late appearing glial cell types. The cell population we studied should therefore be considered as “mixed” but primarily or exclusively neuronal. **Supplementary Table S5** contains the relative expression derived from RNA-Seq read data for selected cell type specific genes normalized to the mean expression for all detected genes and includes data for a few selected primary AD genes.

### Tau related phenotypes

We did not observe genotype specific differences in the levels of phospho-tau, only a redistribution from neural processes to primarily somal regions in edited samples likely related to fewer neuronal processes in dead/dying neurons at late culture times. In contrast, human cell models of FAD iPS cells grown as organoids or FAD genes overexpressed in neural precursor cells differentiated in a 3-D matrix successfully elaborate aspects of tau related phenotypes, even AD-like neurofibrillary tangle formation [25,26]. The 3-D culture format has been suggested to be critical for these tau phenotypes, so the absence of tau pathology in our model could simply be a result of the comparatively small size of the NCs (i.e. ~10-16 μm of thickness and a range of lateral dimensions which varies ~2-3 fold among independent differentiations). An alternative explanation, however is possible. Tau related pathology may be related primarily to non-Aβ dependent APP fragments [77]. These non-Aβ APP fragments are likely present in higher amounts in both the FAD organelle model as well as the FAD overexpression model but not in our edited cells. Consistent with this possibility, cell models constructed with a Down syndrome genotype elaborate both Aβ and tau pathology but similar to these organoid and matrix 3D models, synaptic deficits and ND are not [78]. Increased phosphorylation of tau, but not neurofibrillary tangles, was a feature of degenerating human neurons transplanted into the brain of FAD expressing mice [72]. Clearly additional work is needed to understand the complex relationship between Aβ and tau. It may be significant however that a mouse/human hybrid model and our model suggest that extensive tau pathology is not strictly necessary for progressive Aβ-dependent ND.

### APOE allele type

APOE ε4 and its relationship to AD pathogenesis is complex and incompletely understood, especially with respect to specific brain cell types mediating AD-like phenotypes [79]. APOE allele type involves both Aβ-dependent and independent roles and has been associated with increased neuronal amyloidogenic Aβ production as well as important astrocyte or microglial roles in Aβ removal [80–82]. Since our cultures do not depend on amyloidogenesis and are primarily neuronal, these mechanisms are unlikely to contribute to edit specific phenotypic changes. More likely, APOE ε4 may facilitate neuronal uptake of potentially secreted Aβ42 via endocytosis [83,84]. Alternatively, Aβ may be retained within recycling endosomes since accumulation of aggregated/oligomeric Aβ appears to be initially localized to small putative vesicular compartments and we were unable to detect it in culture media. Either of these possibilities agree with the endolysosomal dysfunction phenotypes we observed and are consistent with an intraneuronal role for Aβ toxicity. While a pathologic role for intraneuronal Aβ was initially contentious, it is now well supported by a variety of evidence, but still lacks a specific mechanism [64].

Neurons under stress are known to increase APOE expression [79] and we did see a modest increase in edited cell lines relative to H9 cells (see **Supplemental Table S5**). CRISPR edited homozygous ε4 human iPS derived neurons have a toxic gain-of-function phenotype which was sufficient to cause ND of GABAergic neurons [85]. This phenotype was human specific, was accompanied by increased amyloidogenesis and also had increased neuronal phospho-tau which was not related to increased Aβ production. Another recent study used independent cell type specific differentiation of edited iPS cells to examine neuronal APOE allele type dependent phenotypes [86] and confirmed that ε4 astrocytes and microglial-like cells can clear extracellular Aβ but also established significant ε4 dependent changes in neuronal gene expression some of which were related to synaptic function which could be relevant to the synaptic deficits we observed.

We also measured very low levels of glial specific marker genes (GFAP, OLIG2 and TREM2 and AIF1; see **Supplementary Table S5**) so we cannot definitively exclude contributions from these non-neuronal cell types without additional observations, however, no positive anti-GFAP staining was detected at any culture stage (data not shown).

### Gene Expression

DEGs that showed a more significant change with greater magnitude in Aβ42 compared to Aβ40 samples suggesting that common pathways may be affected. This agrees well with the exclusive and/or more penetrant phenotypic changes in Aβ42 samples at the time mRNA was isolated for RNA-Seq analysis. There thus appears to be a strong relationship between Aβ-dependent phenotypes and DEGs in these cultures.

Are they also related to AD? We identified a relatively small number of genes compared to extensive AD whole transcriptome expression profiling or other iPS cellular AD models where hundreds to thousands of genes can be differentially expressed [50,87]. This numerical difference could be explained by many factors including intrinsic genetic variance, differences in tissue and cell type sampling, co-morbidities or life style differences not represented in an isogenic culture model. Perhaps more likely, patient material reflects the full spectrum of AD phenotypes, including tau pathology and non-neuronal inflammatory glial responses while our cultures are more likely to represent earlier Aβ-specific putative disease related processes. Despite these considerations, comparison of our data with patient derived RNA-Seq expression data shows some selective overlap. Comparison to an AD vs non-AD study of temporal cortex samples [88] revealed that 88% (82/93) of our DEGs overlap with 51% (42) having the same directional FC. A comparison to a hippocampal study [50] identified 39% (36) genes with overlap and 67% (24) of these had the same directional negative FC (i.e. decreased expression). IPA analysis surprisingly identified “increased neuronal loss” and “decreased memory” two processes with obvious relevance to AD and identified the involvement of the neuroprotective role of THIOP1in AD as a significant disease related pathway. These comparisons and analysis suggest potential relevance or the DEGs to AD which seems more prominent for down regulated genes.

Annotation/enrichment analysis indicates that several of our downregulated DEGs related to cilia, an organelle not usually associated with AD. This suggests that cilia dysfunction may be caused by direct Aβ expression in edited cultures. Five overlapping cilia related genes are also downregulated in AD hippocampus [50] (DAW1, DNAH11, DNAI2, GDA, TEKT1) suggesting that this organelle could also be compromised in patients. Neurons usually contain a primary non-motile cilia [53] believed to function as a major signaling center integrating environmental information through a wide variety of localized G-protein coupled and other types of neuroactive ligand receptors [89]. Notably, the KEGG Neuroactive Ligand Receptor Interaction pathway (see **Supplemental Table S4**), the top scoring pathway we identified by GSEA, suggesting that neuroactive signaling is broadly disrupted in our Aβ42 edited cultures. Ciliary functions have been best studied in sensory neurons which contain specialized types of primary cilia.Interestingly, olfactory neurons have been shown to have Aβ-dependent connectivity defects [90] and be a primary problem in certain types of retinal degeneration [53]. Cerebellar ciliary dysgenesis is prominent in dominant spinocerebellar ataxia type 11 caused by a dominant mutation in tau kinase 2 (TTBK2) [91]. These observations suggest that ciliary dysfunction may thus be intimately related to ND. Neuronal cilia play important roles in neurogenesis, axon guidance, establishment/maintenance of cell polarity, and even synaptic and memory functions [92,93]. Additionally, cilia have a striking similarity to dendritic spines that includes their protein and membrane composition as well as their receptive functions [94]. Our results suggest that disruption of primary cilia may be an important aspect of Aβ-dependent ND which should be examined in future studies.

Future AD therapeutic development may critically depend on identifying specific molecular and cellular mechanisms coupling Aβ to progressive ND [95]. The culture model we describe here may thus be a useful new tool to identify these largely unknown details. This simple neuronal culture model could additionally be a useful tool to identify new therapeutic targets and agents so desperately needed by the growing population of AD patients [96]. While cell culture models may never be able to generate the full complexity of AD disease phenotypes, they are likely to be important for solving many pieces of the puzzle and the exact pathogenic role of Aβ accumulation within or around human neurons seems like an important step forward.

## Acknowledgments

We are grateful to the Sidell-Kagan Foundation for their support of this work. Additional funds were provided by a City of Hope facilities grant for RNA-Seq analysis. Salary support for T.U. was provided by a CIRM Bridges Program Grant (TB1-01185) awarded to Dr. Nicole Bournias. We wish to thank Ms. Tammy Chang for valuable advice and guidance regarding work with human stem cells and Dr. Jessica Kurata for help with the GSEA analysis.

## Availability of materials

The cell lines generated in this study will be made freely available to qualified investigators subject to completing a City of Hope Material Transfer Agreement.

Author contributions
Conceptualization, P.M.S.; Investigation, P.M.S., T.U., M.M., S.S., C.W.; Supervision, P.M.S., J.Y.; Methodology, J.Y., S.S.; Writing-Original Draft P.M.S.; Writing-Review & Editing, PMS, C.W., T.U., M.M., J.Y.; Formal Analysis, C.W.; Visualization, T.U., P.M.S.; Funding Acquisition, P.M.S.
Some of the work in this manuscript formed the basis of a Master of Science degree awarded to T. U. by the Biology Department, California State University San Bernardino.

## References

1. Jack CR, Albert MS, Knopman DS, McKhann GM, Sperling RA, Carrillo MC, et al. Introduction to the recommendations from the National Institute on Aging-Alzheimer’s Association workgroups on diagnostic guidelines for Alzheimer’s disease. Alzheimer’s Dement. Elsevier Ltd; 2011;7: 257–262. doi:10.1016/j.jalz.2011.03.004

2. Hyman BT, Phelps CH, Beach TG, Bigio EH, Cairns NJ, Carrillo MC, et al. National Institute on Aging–Alzheimer’s Association guidelines for the neuropathologic assessment of Alzheimer’s disease. Alzheimer’s Dement. 2012;8: 1–13. doi:10.1016/j.jalz.2011.10.007

3. Cummings JL, Morstorf T, Zhong K. Alzheimer’s disease drug-development pipeline: few candidates, frequent failures. Alzheimers Res Ther. 2014;6: 37. doi:10.1186/alzrt269

4. Sasaguri H, Nilsson P, Hashimoto S, Nagata K, Saito T, De Strooper B, et al. APP mouse models for Alzheimer’s disease preclinical studies. EMBO J. 2017;36: 2473–2487. doi:10.15252/embj.201797397

5. Zahs KR, Ashe KH. “Too much good news” - are Alzheimer mouse models trying to tell us how to prevent, not cure, Alzheimer’s disease? Trends in Neurosciences. 2010. pp. 381–389. doi:10.1016/j.tins.2010.05.004

6. Drummond E, Wisniewski T. Alzheimer’s disease: experimental models and reality. Acta Neuropathol. Springer Berlin Heidelberg; 2017;133: 155–175. doi:10.1007/s00401-016-1662-x

7. Ballatore C, Lee VM-Y, Trojanowski JQ. Tau-mediated neurodegeneration in Alzheimer’s disease and related disorders. Nat Rev Neurosci. 2007;8: 663–672. doi:10.1038/nrn2194

8. Oddo S, Caccamo A, Shepherd JD, Murphy MP, Golde TE, Kayed R, et al. Triple-transgenic model of Alzheimer’s disease with plaques and tangles: intracellular Abeta and synaptic dysfunction. Neuron. 2003/08/05. 2003;39: 409–421. doi:S0896627303004343 [pii]

9. Ghetti B, Oblak AL, Boeve BF, Johnson KA, Dickerson BC, Goedert M. Invited review: Frontotemporal dementia caused by microtubule-associated protein tau gene (MAPT) mutations: a chameleon for neuropathology and neuroimaging. Neuropathol Appl Neurobiol. 2015;41: 24–46. doi:10.1111/nan.12213

10. Iijima-Ando K, Iijima K. Transgenic Drosophila models of Alzheimer’s disease and tauopathies. Brain Struct Funct. 2009;214: 245–262. doi:10.1007/s00429-009-0234-4

11. Abramowski D, Rabe S, Upadhaya AR, Reichwald J, Danner S, Staab D, et al. Transgenic expression of intraneuronal Abeta42 but not Abeta40 leads to cellular Abeta lesions, degeneration, and functional impairment without typical Alzheimer’s disease pathology. J Neurosci. 2012/01/27. 2012;32: 1273–1283. doi:32/4/1273 [pii] 10.1523/JNEUROSCI.4586-11.2012

12. Lewis P, Piper S, Baker M, Onstead L, Murphy M, Hardy J, et al. Expression of BRI-amyloid beta peptide fusion proteins: a novel method for specific high-level expression of amyloid beta peptides. Biochim Biophys Acta. 2001;1537: 58–62. doi:S0925-4439(01)00054-0 [pii]

13. LaFerla FM, Tinkle BT, Bieberich CJ, Haudenschild CC, Jay G. The Alzheimer’s A beta peptide induces neurodegeneration and apoptotic cell death in transgenic mice. Nat Genet. 1995;9: 21–30. Available: http://www.ncbi.nlm.nih.gov/entrez/query.fcgi?cmd=Retrieve&db=PubMed&dopt=Citation&list_uids=7704018

14. McGowan E, Pickford F, Kim J, Onstead L, Eriksen J, Yu C, et al. Aß42 Is Essential for Parenchymal and Vascular Amyloid Deposition in Mice. Neuron. 2005;47: 191–199. doi:10.1016/j.neuron.2005.06.030

15. Iijima K, Liu H-PP, Chiang A-SS, Hearn SA, Konsolaki M, Zhong Y. Dissecting the pathological effects of human Abeta40 and Abeta42 in Drosophila: a potential model for Alzheimer’s disease. Proc Natl Acad Sci U S A. 2004;101: 6623–6628. Available: http://www.ncbi.nlm.nih.gov/entrez/query.fcgi?cmd=Retrieve&db=PubMed&dopt=Citation&list_uids=15069204

16. Ling D, Song H-J, Garza D, Neufeld TP, Salvaterra PM. Abeta42-Induced Neurodegeneration via an Age-Dependent Autophagic-Lysosomal Injury in Drosophila. Cookson MR, editor. PLoS One. 2009;4: e4201. doi:10.1371/journal.pone.0004201

17. Ling D, Magallanes M, Salvaterra PM. Accumulation of Amyloid-Like Aβ 1–42 in AEL (Autophagy–Endosomal–Lysosomal) Vesicles: Potential Implications for Plaque Biogenesis. ASN Neuro. 2014;6: AN20130044. doi:10.1042/AN20130044

18. Selkoe DJ, Hardy J. The amyloid hypothesis of Alzheimer’s disease at 25 years. EMBO Mol Med. 2016;8: 1–14. doi:10.15252/emmm.201606210

19. Musiek ES, Holtzman DM. Three dimensions of the amyloid hypothesis: time, space and “wingmen.” Nat Neurosci. 2015;18: 800–806. doi:10.1038/nn.4018

20. Benilova I, Karran E, De Strooper B. The toxic Aβ oligomer and Alzheimer’s disease: an emperor in need of clothes. Nat Neurosci. Nature Publishing Group; 2012;15: 349–357. doi:10.1038/nn.3028

21. Gordon BA, Blazey TM, Su Y, Hari-Raj A, Dincer A, Flores S, et al. Spatial patterns of neuroimaging biomarker change in individuals from families with autosomal dominant Alzheimer’s disease: a longitudinal study. Lancet Neurol. 2018; 241–250. doi:10.1016/S1474-4422(18)30028-0

22. Mungenast AE, Siegert S, Tsai L-H. Modeling Alzheimer’s disease with human induced pluripotent stem (iPS) cells. Mol Cell Neurosci. Elsevier Inc.; 2016;73: 13–31. doi:10.1016/j.mcn.2015.11.010

23. Israel MA, Yuan SH, Bardy C, Reyna SM, Mu Y, Herrera C, et al. Probing sporadic and familial Alzheimer’s disease using induced pluripotent stem cells. Nature. 2012/01/27. 2012;482: 216–220. doi:10.1038/nature10821

24. Yagi T, Ito D, Okada Y, Akamatsu W, Nihei Y, Yoshizaki T, et al. Modeling familial Alzheimer’s disease with induced pluripotent stem cells. Hum Mol Genet. 2011/09/09. 2011;20: 4530–4539. doi:10.1093/hmg/ddr394

25. Choi SH, Kim YH, Hebisch M, Sliwinski C, Lee S, D’Avanzo C, et al. A three-dimensional human neural cell culture model of Alzheimer’s disease. Nature. Nature Publishing Group; 2014;515: 274–278. doi:10.1038/nature13800

26. Raja WK, Mungenast AE, Lin Y-T, Ko T, Abdurrob F, Seo J, et al. Self-Organizing 3D Human Neural Tissue Derived from Induced Pluripotent Stem Cells Recapitulate Alzheimer’s Disease Phenotypes. Padmanabhan J, editor. PLoS One. 2016;11: e0161969. doi:10.1371/journal.pone.0161969

27. D’Avanzo C, Aronson J, Kim YH, Choi SH, Tanzi RE, Kim DY. Alzheimer’s in 3D culture: Challenges and perspectives. BioEssays. 2015;37: 1139–1148. doi:10.1002/bies.201500063

28. Cermak T, Doyle EL, Christian M, Wang L, Zhang Y, Schmidt C, et al. Efficient design and assembly of custom TALEN and other TAL effector-based constructs for DNA targeting. Nucleic Acids Res. 2011/04/16. 2011;39: e82. doi:gkr218 [pii]10.1093/nar/gkr218

29. Amoroso MW, Croft GF, Williams DJ, O’Keeffe S, Carrasco MA, Davis AR, et al. Accelerated High-Yield Generation of Limb-Innervating Motor Neurons from Human Stem Cells. J Neurosci. 2013;33: 574–586. doi:10.1523/JNEUROSCI.0906-12.2013

30. Schindelin J, Arganda-Carreras I, Frise E, Kaynig V, Longair M, Pietzsch T, et al. Fiji: an open-source platform for biological-image analysis. Nat Methods. Nature Publishing Group, a division of Macmillan Publishers Limited. All Rights Reserved.; 2012;9: 676–682. doi:10.1038/nmeth.2019

31. Genin E, Hannequin D, Wallon D, Sleegers K, Hiltunen M, Combarros O, et al. APOE and Alzheimer disease: a major gene with semi-dominant inheritance. Mol Psychiatry. 2011;16: 903–7. doi:10.1038/mp.2011.52

32. Woodruff G, Young JE, Martinez FJ, Buen F, Gore A, Kinaga J, et al. The Presenilin-1 ?E9 Mutation Results in Reduced ?-Secretase Activity, but Not Total Loss of PS1 Function, in Isogenic Human Stem Cells. Cell Rep. The Authors; 2013;5: 974–985. doi:10.1016/j.celrep.2013.10.018

33. Bergström P, Agholme L, Nazir FH, Satir TM, Toombs J, Wellington H, et al. Amyloid precursor protein expression and processing are differentially regulated during cortical neuron differentiation. Sci Rep. Nature Publishing Group; 2016;6: 29200. doi:10.1038/srep29200

34. Shakes LA, Du H, Wolf HM, Hatcher C, Norford DC, Precht P, et al. Using BAC transgenesis in zebrafish to identify regulatory sequences of the amyloid precursor protein gene in humans. BMC Genomics. 2012;13: 451. doi:10.1186/1471-2164-13-451

35. Davis RP, Costa M, Grandela C, Holland AM, Hatzistavrou T, Micallef SJ, et al. A protocol for removal of antibiotic resistance cassettes from human embryonic stem cells genetically modified by homologous recombination or transgenesis. Nat Protoc. 2008;3: 1550–1558. doi:10.1038/nprot.2008.146

36. Israel MA, Goldstein LS. Capturing Alzheimer’s disease genomes with induced pluripotent stem cells: prospects and challenges. Genome Med. 2011/08/27. 2011;3: 49. doi:10.1186/gm265

37. Muratore CR, Rice HC, Srikanth P, Callahan DG, Shin T, Benjamin LNP, et al. The familial Alzheimer’s disease APPV717I mutation alters APP processing and Tau expression in iPSC-derived neurons. Hum Mol Genet. 2014;23: 3523–3536. doi:10.1093/hmg/ddu064

38. van Helmond Z, Miners JS, Kehoe PG, Love S. Oligomeric Aβ in Alzheimer’s Disease: Relationship to Plaque and Tangle Pathology, APOE Genotype and Cerebral Amyloid Angiopathy. Brain Pathol. 2010;20: 468–480. doi:10.1111/j.1750-3639.2009.00321.x

39. Bharadwaj PR, Dubey AK, Masters CL, Martins RN, Macreadie IG. Abeta aggregation and possible implications in Alzheimer’s disease pathogenesis. J Cell Mol Med. 2009;13: 412–421. Available: http://www.ncbi.nlm.nih.gov/entrez/query.fcgi?cmd=Retrieve&db=PubMed&dopt=Citation&list_uids=19374683

40. Forner S, Baglietto-Vargas D, Martini AC, Trujillo-Estrada L, LaFerla FM. Synaptic Impairment in Alzheimer’s Disease: A Dysregulated Symphony. Trends Neurosci. 2017;40: 347–357. doi:10.1016/j.tins.2017.04.002

41. Nixon RA. Amyloid precursor protein and endosomal-lysosomal dysfunction in Alzheimer’s disease: inseparable partners in a multifactorial disease. FASEB J. 2017;31: 2729–2743. doi:10.1096/fj.201700359

42. Nilsson P, Loganathan K, Sekiguchi M, Matsuba Y, Hui K, Tsubuki S, et al. Aß Secretion and Plaque Formation Depend on Autophagy. Cell Rep. The Authors; 2013;7: 1–9. doi:10.1016/j.celrep.2013.08.042

43. Cataldo AM, Petanceska S, Terio NB, Peterhoff CM, Durham R, Mercken M, et al. Abeta localization in abnormal endosomes: association with earliest Abeta elevations in AD and Down syndrome. Neurobiol Aging. 2004/10/07. 2004;25: 1263–1272. doi:10.1016/j.neurobiolaging.2004.02.027

44. Poteryaev D, Datta S, Ackema K, Zerial M, Spang A. Identification of the switch in early-to-late endosome transition. Cell. Elsevier Ltd; 2010;141: 497–508. doi:10.1016/j.cell.2010.03.011

45. Schluter OM. A Complete Genetic Analysis of Neuronal Rab3 Function. J Neurosci. 2004;24: 6629–6637. doi:10.1523/JNEUROSCI.1610-04.2004

46. Boland B, Kumar A, Lee S, Platt FM, Wegiel J, Yu WH, et al. Autophagy Induction and Autophagosome Clearance in Neurons: Relationship to Autophagic Pathology in Alzheimer’s Disease. J Neurosci. 2008;28: 6926–6937. doi:10.1523/jneurosci.0800-08.2008

47. Haass C, Kaether C, Thinakaran G, Sisodia S. Trafficking and proteolytic processing of APP. Cold Spring Harb Perspect Med. 2012;2: a006270. doi:10.1101/cshperspect.a006270

48. Grueninger F, Bohrmann B, Czech C, Ballard TM, Frey JR, Weidensteiner C, et al. Phosphorylation of Tau at S422 is enhanced by Aβ in TauPS2APP triple transgenic mice. Neurobiol Dis. Elsevier Inc.; 2010;37: 294–306. doi:10.1016/j.nbd.2009.09.004

49. Corder AEH, Saunders AM, Strittmatter WJ, Schmechel DE, Gaskell PC, Small W, et al. Gene Dose of Apolipoprotein E Type 4 Allele and the Risk of Alzheimer’s Disease in Late Onset Families Published by?: American Association for the Advancement of Science Stable URL?: http://www.jstor.org/stable/2882127. 2008;261: p921–923.

50. Annese A, Manzari C, Lionetti C, Picardi E, Horner DS, Chiara M, et al. Whole transcriptome profiling of Late-Onset Alzheimer’s Disease patients provides insights into the molecular changes involved in the disease. Sci Rep. 2018;8: 4282. doi:10.1038/s41598-018-22701-2

51. Blalock EM, Geddes JW, Chen KC, Porter NM, Markesbery WR, Landfield PW. Incipient Alzheimer’s disease: Microarray correlation analyses reveal major transcriptional and tumor suppressor responses. Proc Natl Acad Sci. 2004;101: 2173–2178. doi:10.1073/pnas.0308512100

52. Zhang B, Gaiteri C, Bodea L-G, Wang Z, McElwee J, Podtelezhnikov AA, et al. Integrated systems approach identifies genetic nodes and networks in late-onset Alzheimer’s disease. Cell. Elsevier Inc.; 2013;153: 707–20. doi:10.1016/j.cell.2013.03.030

53. Guemez-Gamboa A, Coufal NG, Gleeson JG. Primary Cilia in the Developing and Mature Brain. Neuron. Elsevier; 2014;82: 511–521. doi:10.1016/j.neuron.2014.04.024

54. Yamin R, Malgeri EG, Sloane J a, Mcgraw WT, Abraham CR. Metalloendopeptidase EC 3. 4. 24. 15 Is Necessary for Alzheimer’s Amyloid-? Peptide Degradation *. Biochemistry. 1999;

55. Pollio G, Hoozemans JJM, Andersen CA, Roncarati R, Rosi MC, van Haastert ES, et al. Increased expression of the oligopeptidase THOP1 is a neuroprotective response to Aß toxicity. Neurobiol Dis. 2008;31: 145–158. doi:10.1016/j.nbd.2008.04.004

56. Saghatelian A, Jessani N, Joseph A, Humphrey M, Cravatt BF. Activity-based probes for the proteomic profiling of metalloproteases. Proc Natl Acad Sci. 2004;101: 10000–10005. doi:10.1073/pnas.0402784101

57. Abraham CR, Selkoe DJ, Potter H. Immunochemical identification of the serine protease inhibitor a1-antichymotrypsin in the brain amyloid deposits of Alzheimer’s disease. Cell. 1988;52: 487–501. doi:10.1016/0092-8674(88)90462-X

58. Freude K, Pires C, Hyttel P, Hall V. Induced Pluripotent Stem Cells Derived from Alzheimer’s Disease Patients: The Promise, the Hope and the Path Ahead. J Clin Med. 2014;3: 1402–1436. doi:10.3390/jcm3041402

59. Young JE, Goldstein LSB. Alzheimer’s disease in a dish: promises and challenges of human stem cell models. Hum Mol Genet. 2012;21: R82–R89. doi:10.1093/hmg/dds319

60. Arber C, Lovejoy C, Wray S. Stem cell models of Alzheimer’s disease: progress and challenges. Alzheimers Res Ther. Alzheimer’s Research & Therapy; 2017;9: 42. doi:10.1186/s13195-017-0268-4

61. Müller UC, Deller T, Korte M. Not just amyloid: physiological functions of the amyloid precursor protein family. Nat Rev Neurosci. Nature Publishing Group; 2017;18: 281–298. doi:10.1038/nrn.2017.29

62. Tanzi RE. The genetics of Alzheimer disease. Cold Spring Harb Perspect Med. 2012;2: 1– 10. doi:10.1101/cshperspect.a006296

63. Herrup K. The case for rejecting the amyloid cascade hypothesis. Nat Neurosci. 2015;18: 794–799. doi:10.1038/nn.4017

64. Gouras GK, Tampellini D, Takahashi RH, Capetillo-Zarate E. Intraneuronal ß-amyloid accumulation and synapse pathology in Alzheimer’s disease. Acta Neuropathol. 2010;119: 523–541. doi:10.1007/s00401-010-0679-9

65. Lopez-Toledano MA. Neurogenic Effect of -Amyloid Peptide in the Development of Neural Stem Cells. J Neurosci. 2004;24: 5439–5444. doi:10.1523/JNEUROSCI.0974-04.2004

66. Hu X, Crick SL, Bu G, Frieden C, Pappu R V, Lee J-M. Amyloid seeds formed by cellular uptake, concentration, and aggregation of the amyloid-beta peptide. Proc Natl Acad Sci U S A. 2009;106: 20324–9. doi:10.1073/pnas.0911281106

67. Gouras GK, Almeida CG, Takahashi RH. Intraneuronal Abeta accumulation and origin of plaques in Alzheimer’s disease. Neurobiol Aging. 2005;26: 1235–1244. Available: http://www.ncbi.nlm.nih.gov/entrez/query.fcgi?cmd=Retrieve&db=PubMed&dopt=Citation&list_uids=16023263

68. Galluzzi L, Vitale I, Aaronson SA, Abrams JM, Adam D, Agostinis P, et al. Molecular mechanisms of cell death: recommendations of the Nomenclature Committee on Cell Death 2018. Cell Death Differ. 2018;25: 486–541. doi:10.1038/s41418-017-0012-4

69. Ghavami S, Shojaei S, Yeganeh B, Ande SR, Jangamreddy JR, Mehrpour M, et al. Autophagy and apoptosis dysfunction in neurodegenerative disorders. Prog Neurobiol. Elsevier Ltd; 2014;112: 24–49. doi:10.1016/j.pneurobio.2013.10.004

70. Zhang Y, McLaughlin R, Goodyer C, LeBlanc A. Selective cytotoxicity of intracellular amyloid ß peptide1-42through p53 and Bax in cultured primary human neurons. J Cell Biol. 2002;156: 519–529. doi:10.1083/jcb.200110119

71. Moore BD, Martin J, de Mena L, Sanchez J, Cruz PE, Ceballos-Diaz C, et al. Short Aß peptides attenuate Aß42 toxicity in vivo. J Exp Med. 2018;215: 283–301. doi:10.1084/jem.20170600

72. Espuny-Camacho I, Arranz AM, Fiers M, Snellinx A, Ando K, Munck S, et al. Hallmarks of Alzheimer’s Disease in Stem-Cell-Derived Human Neurons Transplanted into Mouse Brain. Neuron. 2017;93: 1066–1081.e8. doi:10.1016/j.neuron.2017.02.001

73. Haass C, Selkoe DJ. Soluble protein oligomers in neurodegeneration: lessons from the Alzheimer’s amyloid beta-peptide. Nat Rev Mol Cell Biol. 2007;8: 101–12. doi:10.1038/nrm2101

74. Wicklund L, Leão RN, Strömberg A-M, Mousavi M, Hovatta O, Nordberg A, et al. ?-Amyloid 1-42 Oligomers Impair Function of Human Embryonic Stem Cell-Derived Forebrain Cholinergic Neurons. PLoS One. 2010;5: e15600. doi:10.1371/journal.pone.0015600

75. Engel M, Do-Ha D, Muñoz SS, Ooi L. Common pitfalls of stem cell differentiation: a guide to improving protocols for neurodegenerative disease models and research. Cell Mol Life Sci. 2016;73: 3693–3709. doi:10.1007/s00018-016-2265-3

76. Muratore CR, Zhou C, Liao M, Fernandez MA, Taylor WM, Lagomarsino VN, et al. Cell-type Dependent Alzheimer’s Disease Phenotypes: Probing the Biology of Selective Neuronal Vulnerability. Stem Cell Reports. ElsevierCompany.; 2017;9: 1868–1884. doi:10.1016/j.stemcr.2017.10.015

77. Moore S, Evans LDB, Andersson T, Portelius E, Smith J, Dias TB, et al. APP Metabolism Regulates Tau Proteostasis in Human Cerebral Cortex Neurons. Cell Rep. Elsevier; 2015;11: 689–696. doi:10.1016/j.celrep.2015.03.068

78. Shi Y, Kirwan P, Smith J, MacLean G, Orkin SH, Livesey FJ. A human stem cell model of early Alzheimer’s disease pathology in Down syndrome. Sci Transl Med. 2012/02/22. 2012;4: 124ra29–124ra29. doi:10.1126/scitranslmed.3003771

79. Mahley RW, Huang Y. Apolipoprotein E Sets the Stage: Response to Injury Triggers Neuropathology. Neuron. 2012;76: 871–885. doi:10.1016/j.neuron.2012.11.020

80. Efthymiou AG, Goate AM. Late onset Alzheimer’s disease genetics implicates microglial pathways in disease risk. Mol Neurodegener. Molecular Neurodegeneration; 2017;12: 43. doi:10.1186/s13024-017-0184-x

81. Wyss-Coray T, Loike JD, Brionne TC, Lu E, Anankov R, Yan F, et al. Adult mouse astrocytes degrade amyloid-β in vitro and in situ. Nat Med. 2003;9: 453–457. doi:10.1038/nm838

82. Huang Y. Aß-independent roles of apolipoprotein E4 in the pathogenesis of Alzheimer’s disease. Trends Mol Med. Elsevier Ltd; 2010;16: 287–294. doi:10.1016/j.molmed.2010.04.004

83. Dafnis I, Stratikos E, Tzinia A, Tsilibary EC, Zannis VI, Chroni A. An apolipoprotein E4 fragment can promote intracellular accumulation of amyloid peptide beta 42. J Neurochem. 2010;115: 873–884. doi:10.1111/j.1471-4159.2010.06756.x

84. Kanekiyo T, Cirrito JR, Liu C-C, Shinohara M, Li J, Schuler DR, et al. Neuronal Clearance of Amyloid-by Endocytic Receptor LRP1. J Neurosci. 2013;33: 19276–19283. doi:10.1523/JNEUROSCI.3487-13.2013

85. Wang C, Najm R, Xu Q, Jeong DE, Walker D, Balestra ME, et al. Gain of toxic apolipoprotein E4 effects in human iPSC-derived neurons is ameliorated by a small-molecule structure corrector article. Nat Med. Springer US; 2018;24: 647–657. doi:10.1038/s41591-018-0004-z

86. Lin Y-T, Seo J, Gao F, Feldman HM, Wen H-L, Penney J, et al. APOE4 Causes Widespread Molecular and Cellular Alterations Associated with Alzheimer’s Disease Phenotypes in Human iPSC-Derived Brain Cell Types. Neuron. Elsevier Inc.; 2018; 1–14. doi:10.1016/j.neuron.2018.05.008

87. Castillo E, Leon J, Mazzei G, Abolhassani N, Haruyama N, Saito T, et al. Comparative profiling of cortical gene expression in Alzheimer’s disease patients and mouse models demonstrates a link between amyloidosis and neuroinflammation. Sci Rep. 2017;7: 17762. doi:10.1038/s41598-017-17999-3

88. Allen M, Carrasquillo MM, Funk C, Heavner BD, Zou F, Younkin CS, et al. Human whole genome genotype and transcriptome data for Alzheimer’s and other neurodegenerative diseases. Sci Data. 2016;3: 160089. doi:10.1038/sdata.2016.89

89. Berbari NF, O’Connor AK, Haycraft CJ, Yoder BK. The Primary Cilium as a Complex Signaling Center. Curr Biol. Elsevier Ltd; 2009;19: R526–R535. doi:10.1016/j.cub.2009.05.025

90. Cao L, Schrank BR, Rodriguez S, Benz EG, Moulia TW, Rickenbacher GT, et al. Aß alters the connectivity of olfactory neurons in the absence of amyloid plaques in vivo. Nat Commun. Nature Publishing Group; 2012;3: 1009. doi:10.1038/ncomms2013

91. Houlden H, Johnson J, Gardner-Thorpe C, Lashley T, Hernandez D, Worth P, et al. Mutations in TTBK2, encoding a kinase implicated in tau phosphorylation, segregate with spinocerebellar ataxia type 11. Nat Genet. 2007;39: 1434–1436. doi:10.1038/ng.2007.43

92. Lee JH, Gleeson JG. The role of primary cilia in neuronal function. Neurobiol Dis. Elsevier Inc.; 2010;38: 167–172. doi:10.1016/j.nbd.2009.12.022

93. Berbari NF, Malarkey EB, Yazdi SMZR, McNair AD, Kippe JM, Croyle MJ, et al. Hippocampal and Cortical Primary Cilia Are Required for Aversive Memory in Mice. Kavushansky A, editor. PLoS One. 2014;9: e106576. doi:10.1371/journal.pone.0106576

94. Nechipurenko I V., Doroquez DB, Sengupta P. Primary cilia and dendritic spines: Different but similar signaling compartments. Mol Cells. 2013;36: 288–303. doi:10.1007/s10059-013-0246-z

95. Cummings J, Aisen PS, DuBois B, Frölich L, Jack CR, Jones RW, et al. Drug development in Alzheimer’s disease: the path to 2025. Alzheimer’s Res Ther. Alzheimer’s Research & Therapy; 2016;8: 39. doi:10.1186/s13195-016-0207-9

96. Katsnelson A, De Strooper B, Zoghbi HY. Neurodegeneration: From cellular concepts to clinical applications. Sci Transl Med. 2016;8: 1–6. doi:10.1126/scitranslmed.aal2074

